# Data-driven probabilistic definition of the low energy conformational states of protein residues

**DOI:** 10.1101/2023.07.24.550386

**Authors:** Jose Gavalda-Garcia, David Bickel, Joel Roca-Martinez, Daniele Raimondi, Gabriele Orlando, Wim Vranken

## Abstract

Protein dynamics and related conformational changes are essential for their function but difficult to characterise and interpret. Amino acids in a protein behave according to their local energy landscape, which is determined by their local structural context and environmental conditions. The lowest energy state for a given residue can correspond to sharply defined conformations, *e*.*g*., in a stable helix, or can cover a wide range of conformations, *e*.*g*., in intrinsically disordered regions. A good definition of such low energy states is therefore important to describe the behavior of a residue and how it changes with its environment. We propose a data-driven probabilistic definition of six low energy conformational states typically accessible for amino acid residues in proteins. This definition is based on solution NMR information of 1,322 proteins through a combined analysis of structure ensembles with interpreted chemical shifts. We further introduce a conformational state variability parameter that captures, based on an ensemble of protein structures from molecular dynamics or other methods, how often a residue moves between these conformational states. The approach enables a different perspective on the local conformational behavior of proteins that is complementary to their static interpretation from single structure models.

## Introduction

The dynamics and related conformational changes of proteins are often essential for their function, but are difficult to characterise and interpret. A key challenge is that this behavior is typically not encompassed by the available data on protein conformation at atomic resolution, which mostly derive from static representations of the most stable protein conformations in crystalline state(s). The recent AlphaFold2 protein fold predictions are trained on such data and therefore essentially capture those static crystalline states. The dynamic and environmentally determined behavior that proteins can exhibit in solution is, on the other hand, exemplified by intrinsically disordered regions (IDRs) and proteins (IDPs). These do not have a stable conformation but still perform essential functions in physiological conditions, often by temporarily acquiring a fold, for example when interacting with a binding partner [1]. Changes in conformation are also necessary for prevalent allosteric mechanisms in folded proteins [2], with even entire fold changes observed for the same protein [3].

To study a protein’s conformational variability a variety of techniques can be applied, each with their own advantages and challenges. X-ray crystallography provides high-resolution structures but its ability to capture conformational changes is limited. Even though B-factors can be linked to protein flexibility and rigidity [4], the non-native conditions within crystals do not provide a realistic picture of dynamic protein behavior, even at room temperature [5]. Single particle cryo-electron microscopy, in contrast, allows the characterisation of proteins in near-native states. The resolution of the resulting structures directly correlates to the conformational diversity of the protein, with the possibility to classify heterogenous structural ensembles into different conformations [6]. Nuclear magnetic resonance (NMR) provides in-solution information on dynamics of proteins at physiological conditions, from *S*^2^ order parameters determined by relaxation experiments, which capture fast ps-ns timescale movements, to chemical shifts, which cover up to low ms timescale movements [7]. The observations in NMR are typically averaged over all the protein molecules present in the sample, however. Thus, if a protein adopts multiple conformations between which it rapidly interchanges, the observed NMR data will show the linear average of a given parameter over the populations of these conformations. Hydrogen-deuterium exchange mass spectrometry (HDX-MS) provides residue- to peptide-resolution information on conformational changes in the ms-s time scale. Similar to NMR, the outcomes are averaged so that individual conformational populations cannot be resolved. Still, the technology is powerful in its ability to identify dynamic regions in proteins [8]. Single- molecule Forster resonance energy transfer (smFRET) provides distance information on individual conformations in dynamic mixtures. For FRET experiments the proteins need to be dye labelled. These dyes are typically linked by highly flexible linkers to proteins, lowering the accuracy of the measurements [9]. While all of these techniques allow the observation of the conformational diversity of proteins, there is no computational framework available to describe varying propensities towards different experimentally observed conformational states in variable protein regions.

A major challenge in describing the conformational variability of proteins is that their conformational space is extremely high-dimensional. Each amino acid residue in the protein backbone that is not fixed to a single conformation adds to the overall conformational complexity of the protein. Conversely, this provides the opportunity to describe the conformational diversity of a protein in terms of the local conformational freedom of the individual residues. A residue’s conformational preferences are determined by its structural context, interactions with other molecules and environmental conditions such as pH or ionic strength, all of which shape its local energy surface. The lowest energy state for a given residue can then correspond to very sharply defined conformations, for example when the residue is part of a stable helix, or can cover a wide range of conformations, for example for residues in intrinsically disordered regions. Amino acid residues that do change conformation under native conditions can therefore, by definition, not be assigned to a specific secondary structure. They should rather be interpreted in probabilistic terms within the secondary structure space where they transition. This requires a definition of low energy conformational states that residues typically adopt in proteins, as well as a description of the conformational variability within these low energy conformational states. To derive such low energy states, ideally, large scale experimental data should be employed to extract statistically relevant behaviors.

Information about the behavior of amino acids in proteins can be estimated from chemical shifts, the most readily publicly available NMR parameter from the Biological Magnetic Resonance Data Bank (BMRB) [10]. This has resulted in methods to estimate the backbone and side-chain dynamics of amino acids [11, 12], secondary structure populations [13], and a combined residue-independent metric based on unsupervised machine learning [14]. Such parameters summarize the statistically averaged behavior of residues of proteins in solution, so reflecting their lowest energy state in the employed experimental conditions. NMR structure ensembles are also calculated from NMR observables, notably interatomic distances from the nuclear Overhauser effect (NOE), dihedral angle ranges from dipolar couplings and chemical shifts and bond orientations from residual dipolar couplings (RDCs). Typically, time-averaged NMR data are used to calculate a single NMR model at a time, so resulting in information clashes where mutually incompatible conformations are present in the protein [15]. NMR structure ensembles are typically collections of such models, therefore residues that adopt a single well-defined conformation in solution, with all experimental data consistent, will tend to have the same consistent structural definition in all models of the NMR ensemble. For residues that adopt multiple distinct conformations in large enough populations, on the other hand, the observed experimental data will be incompatible within the same structure, so resulting in variable structure definitions in the models of the NMR ensemble. The concept of using such information from NMR ensembles was already successfully employed in for example the Espritz-NMR disorder predictor [16].

However, capturing the inherently probabilistic information from protein structure models that encompass multiple conformations is difficult. To date, no method is available that can describe varying propensities towards different conformational states in highly variable regions. We here propose a data-driven probabilistic definition of six low energy conformational states that are typically accessible for amino acid residues in proteins. This definition is based on solution NMR information for 1,322 proteins, through a combined analysis of structure ensembles with interpreted chemical shifts. Each conformational state corresponds to a specific region in the backbone dihedral space (Fig. 1). Within these regions, residues have the freedom to adopt various conformations without encountering significant energy barriers. We further introduce a conformational state variability parameter that captures how often a residue moves between these conformational states. Such transitions would indicate changes in the local environment of the residues, so resulting in a change of their low energy state. The approach therefore enables a different perspective on the conformational behavior of proteins that is complementary to their static interpretation from single structure models. To demonstrate the effectiveness of our method, we apply it to a set of molecular dynamics (MD) simulations consisting of 113 proteins that exhibit diverse behaviors ranging from structurally rigid to highly dynamic disordered proteins. Additionally, we apply our method to large ensembles of selected intrinsically disordered proteins (IDPs) to showcase its ability to identify regions of relative order within otherwise disordered proteins.We implemented the approach as a python package, named Constava, that is available on PyPI: https://pypi.org/project/constava/.

**Figure 1:**
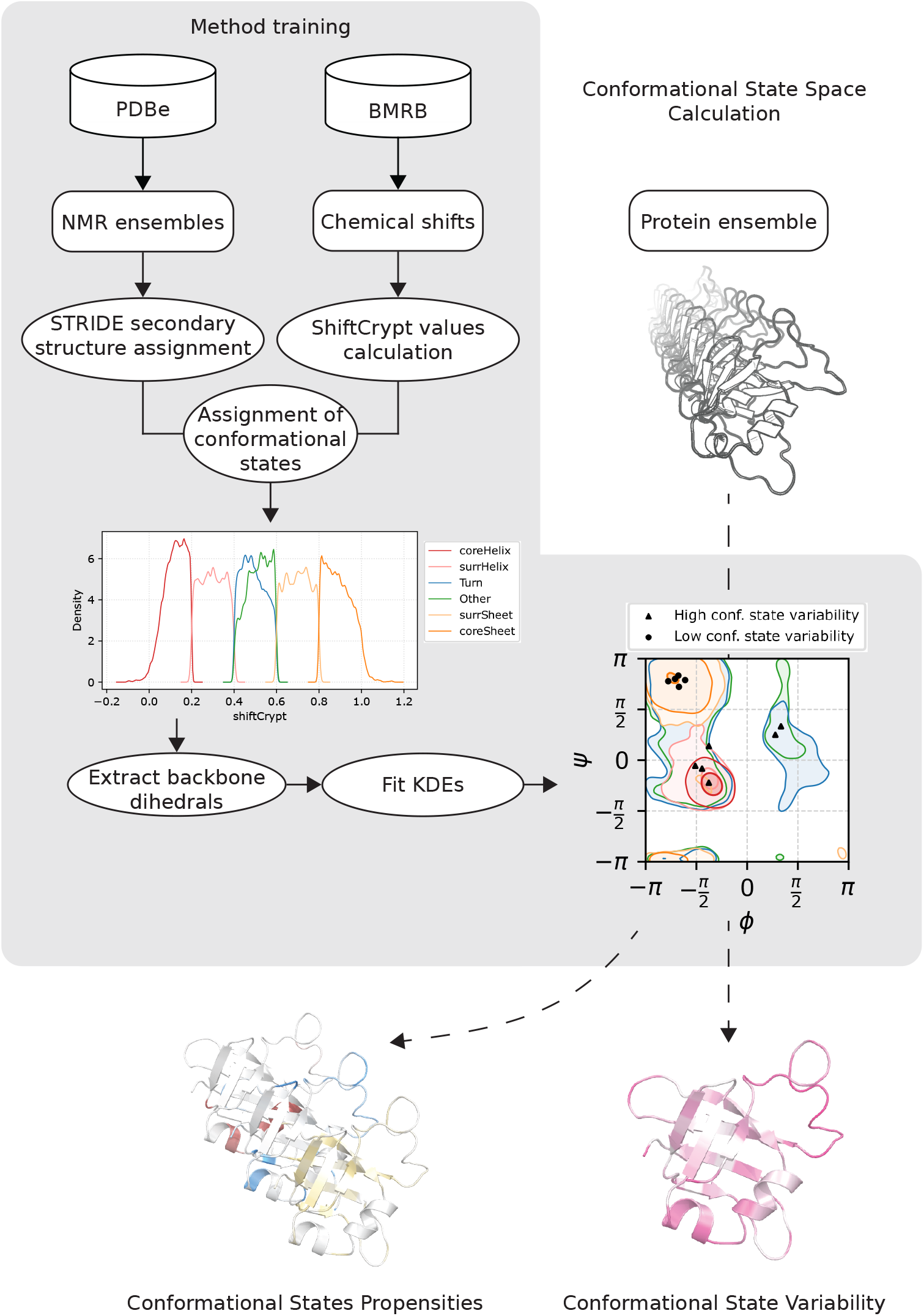
Visual representation of the KDE training workflow and the calculation of confor- mational state propensities and conformational state variability. The training of the models (grey background region) employs NMR ensembles from the PDBe and chemical shift information from the BMRB. The NMR ensembles are used to obtain secondary structure assignments of each residue. The chemical shifts are employed to calculate the ShiftCrypt values. Both data sources are combined to assign the residues to a conformational state. Once the conformational state assignment of these residues is completed, their backbone dihedrals are extracted and used to fit six KDEs, one per conformational state. With these trained KDEs the conformational states probabilities as well as the conformational state variability can be calculated (dashed arrows).

## Methods

An overview of the connection between the Methods sections below is provided in Fig. 1.

### Generation of probabilistic model of conformational states

#### Data collection

Chemical shift data were collected from the Biological Magnetic Resonance Data Bank (BMRB) [10], using previously described criteria [17]. Summarized, only entries between reported pH 5–7, temperature 293–313K were selected for which chemical shift data were available for ^1^H, ^13^C and ^15^N for at least half the residues in the sequence. Entries with samples containing agents that strongly influence protein behaviour (*e*.*g*., TFE, SDS, *etc*.; for complete list consult Table S1 in [17]) were excluded, and chemical shift re-referencing was performed with VASCO [18]. Only BMRB entries with corresponding NMR ensembles of at least 10 models were retained, resulting in 1,414 entries totaling 147,116 residues. STRIDE was applied to all individual models of the NMR ensemble [19, 20]. STRIDE uses structural information to assign secondary structure labels to each residue and thereby provides a categorical description of the conformation of that residue (for each model in the NMR ensemble). ShiftCrypt [14] was applied to the chemical shifts to obtain a single value per residue in the range 0 to 1 that relates to that residue’s behavior in relation to secondary structure propensity and backbone dynamics. ShiftCrypt values near the extremes point towards residues that adopt rigid helix (near 0) or sheet (near 1) conformations, while deviations towards the center indicate increasingly dynamic residues with decreasing degrees of helix/sheet propensity. The data are available in Zenodo: https://doi.org/10.5281/zenodo.10371447.

#### Conformational state definition

Based on the ShiftCrypt values and STRIDE [19, 20] secondary structure assignments of residues in NMR ensembles, we define six conformational states. We then annotated all residues in the NMR ensembles according to these conformational states, resulting in 62,125 annotated residues from 1,322 protein ensembles. To obtain structural insights on conformational space that residues in each state cover, we extracted the backbone dihedral angles *ϕ* and *ψ* for all residues in a given category. To avoid the over-/underrepresentation of individual structures based on the number of models deposited in their NMR ensembles, we randomly selected 5 models for each structural ensemble assigned to a given BMRB ID. This yielded a set of (*ϕ, ψ*)-pairs for each of the conformational states.

- *Core helix* : Residues that exclusively adopt helical conformation in all models of their associated NMR ensemble with shiftCrypt values ≤ 0.2; *N* = 93, 957 residues
- *Surrounding helix* : Residues that adopt helical conformation in the majority of models of their associated NMR ensemble with shiftCrypt values in the range (0.2, 0.4]; *N* = 8, 180 residues
- *Core sheet* : Residues that exclusively adopt extended conformation in all models of their associated NMR ensemble with shiftCrypt values ≥ 0.8; *N* = 47, 280 residues
- *Surrounding sheet* : Residues that adopt extended conformation in the majority of models of their associated NMR ensemble with shiftCrypt values in the range [0.6, 0.8); *N* = 11, 280residues
- *Turn*: Residues that adopt turn conformation in the majority of models of their associated NMR ensemble with shiftCrypt values in the range (0.4, 0.6); *N* = 75, 377 residues
- *Other* : Residues that adopt coil conformation in the majority of models of their associated NMR ensemble with shiftCrypt values in the range (0.4, 0.6); *N* = 74, 542 residues

Gaussian kernel density estimators were then built for each conformational state using the scikit-learn python package with bandwidth for the Gaussians equal to 0.13 radians. They represent how likely it is that a residue in a conformational state (as defined above) occupies a certain region in the backbone dihedral space as observed in the NMR ensemble. The robustness of these probabilistic models is assessed in Supplementary Fig. 1.

### Inference of conformational state propensities

Given a conformational ensemble, the kernel density estimates can then be used to infer the conformational state propensities for each residue. To do so, (*ϕ, ψ*)-angle pairs are extracted from the conformational ensemble and divided in *M* samples of *N* (*ϕ, ψ*)-pairs. This can be done by bootstrapping *N* data points *M* times, or by using a sliding window of size *N* . For each of these samples the evidence is calculated by evaluating the probability density functions (PDF) at the given (*ϕ, ψ*)- coordinates (equation (1)). Then, the likelihood for any state is its evidence relative to all other evidence (equation (2)). We generally report the average over all *M* samples obtained from the ensemble.

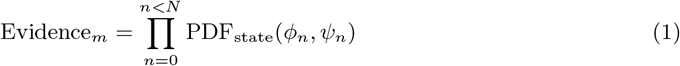

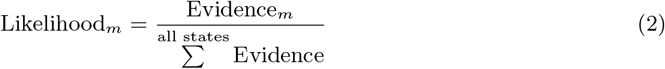

Notably, the sample size *N* chosen to calculate the evidence strongly impacts the certainty with which conformational states are inferred. For small sample sizes (*N* ≤ 5), the method captures likelihoods that reflect the local backbone dynamics and conformational ambiguity, while for larger sample sizes (*N* > 20) the likelihoods result in a categorical assignment of a preferred conformational state for most residues (see Supplementary Fig. 2). In general, we suggest a value of *N* = 3 to obtain information of the conformational state variability (see below), and a value of *N* = 25 to extract preferred conformational states.

**Figure 2:**
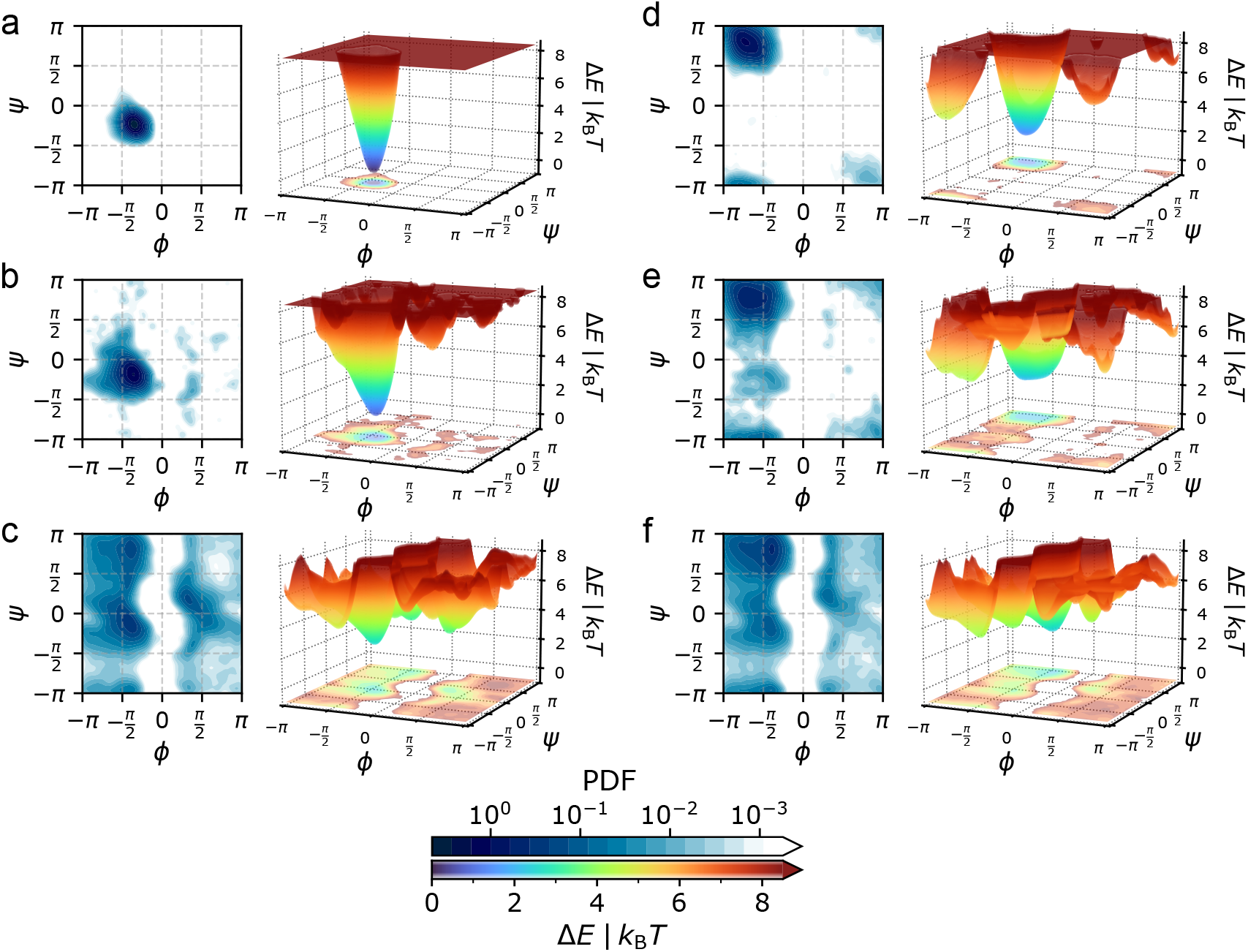
Definition of the six conformational states in (*ϕ, ψ*)-space. Each panel shows a conformational state represented by a continuous probability density function on the left, and the derived potential energy surfaces in the (*ϕ, ψ*)-space on the right. The conformational states are shown for: a) *Core helix*, b) *Surrounding helix*, c) *Turn*, d) *Core sheet*, e) *Surrounding sheet*, and f) *Other*. The potential energy surfaces illustrate how, *e*.*g*., a *Core helix* residue is conformationally restricted by high energy barriers, while *Turn* residues can adopt a wide range of backbone conformations without having to overcome such high energy barriers. This is further exemplified by the 2D projection in Supplementary Fig. 12.

**Figure 3:**
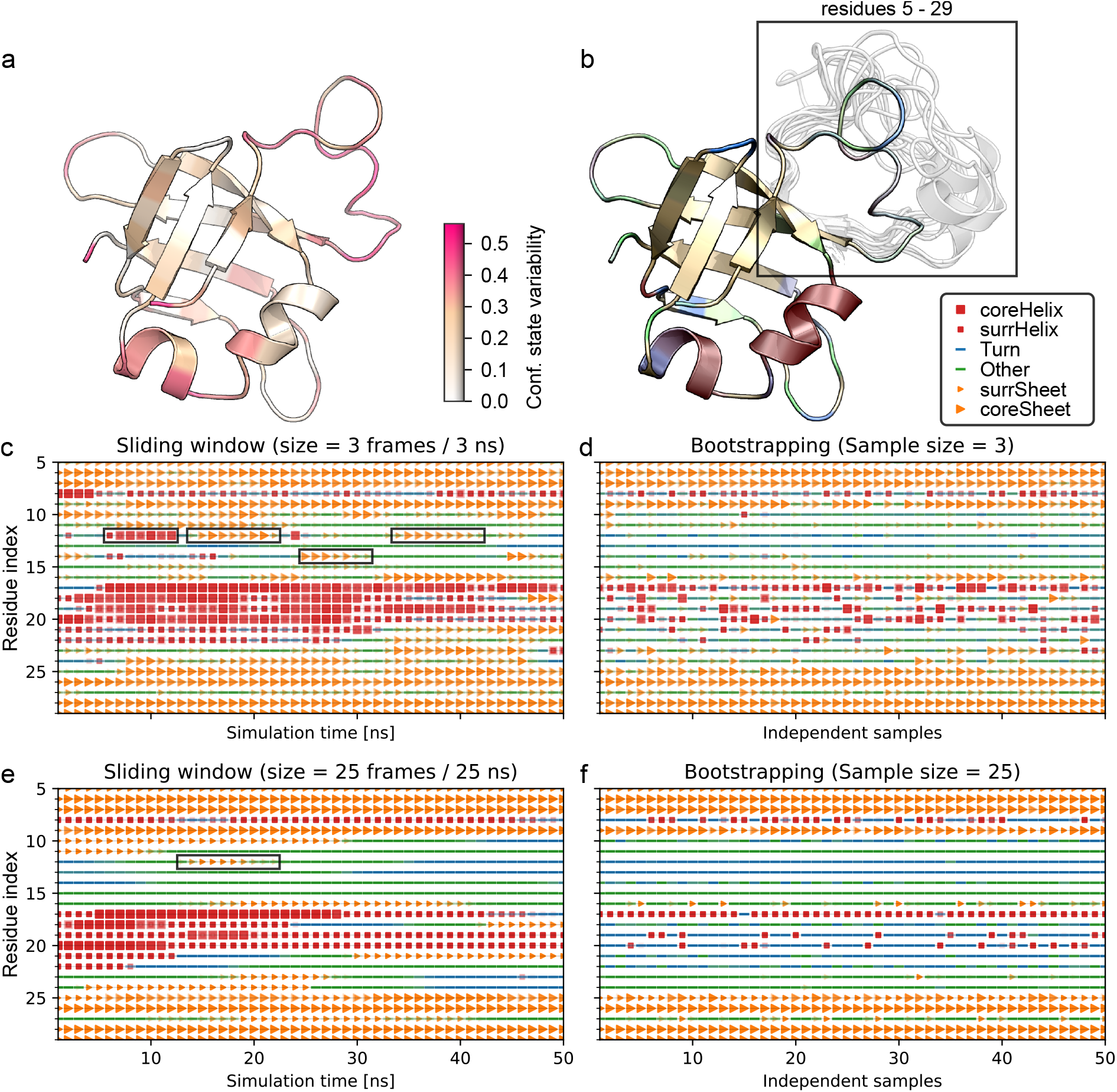
Analysis of MD simulation ensembles with Constava. The method was applied on the conformational ensemble derived from 100 ns of MD simulation on *E. coli* ribosomal protein L25 (PDB ID: 1b75). a) Conformational state variability mapped on the structure. Regions of high variability are the loop region as well as the C-terminal end of the helical region, which switches between helical and turn-like states. b) Conformational states propensities mapped on the structure. The rectangle highlights residues 5-29, where the largest conformational changes occur. c) Conformational states as a time series along the simulation, using a sliding window with *N* = 3 (here 3 ns). Black boxes highlight transient conformational states apparent at this time-resolution. d) Conformational states obtained from bootstrapping (*N* = 3). Adjacent samples are not related. Transient states are sometimes detected, but are underrepresented in comparison to the sliding-window method. e) Conformational states as a time series along the simulation, using a sliding window with *N* = 25 (here 25 ns). Black boxes highlight transient conformational states apparent at this time-resolution. Notably, fewer transient states are detected compared to the smaller window size (panel c). f) Conformational states obtained from bootstrapping (*N* = 25). With increasing sample size the likelihood to detect low populated transient conformational states further diminishes, and the detected conformational states increasingly converge on unique solutions.

### Definition of conformational state variability

In a conformational ensemble a residue may not only undergo conformational changes (*i*.*e*., variation of its *ϕ*- and *ψ*-angles captured by the conformational state KDEs), but may change its conformational preference altogether. We label this as conformational state variability, expressing the ability of a residue to switch back and forth between conformational states. Note that this does not directly relate to changes of the *ϕ*- and *ψ*-angles but captures the ability of a residue to exist in multiple conformational states, taking into account the structural variation that would be expected for any residue in that given conformational state.

To calculate the conformational state variability of a given residue in an ensemble we calculate the root mean square distance from the average state across all *M* samples (equation (3)). Here, *L*_*m*_ is the likelihood of one sample to belong to a conformational state as described in equation (2) and ⟨*L*_*m*_⟩ is the average likelihood across all samples.

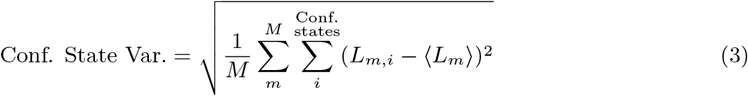

### Molecular dynamics simulations

Reference molecular dynamics (MD) simulations were performed for 113 proteins in order to validate the method proposed in this work and to identify parameters that allow the study of different aspects of conformational state variability. The simulated proteins cover a wide range of proteins, from disordered to partially ordered to fully ordered. The disordered and partially ordered entries were selected from MobiDB [21] based on their coil percentage, using as criteria more than 90% coil (disordered) and approximately 50% coil (partially ordered), respectively. Only entries with an annotation coverage higher than 90% were included. A diverse set of rigid structures was also defined by selecting proteins with different architectures and topologies based on their CATH code [22]. All the selected proteins had structures available that were experimentally determined by NMR and had an associated entry on BMRB with extensive NMR chemical shift data available (^13^C, ^15^N, and ^1^H). The full protein list with the selected PDB codes that were used as starting conformations is available as supplementary information (see Supplementary Data 1).

The MD simulations were prepared and performed with GROMACS [23] using the CHARMM36m force field [24]. All 113 proteins were subjected to the same protocol to yield an internally consistent data set. Starting from the PDB structure (1^st^ model of an NMR ensemble), water molecules and heteroatoms were removed. The systems were solvated using TIP3P water [25] in a rhombic dodecahedron simulation box and Na^+^ or Cl^-^ ions were added to neutralize the systems. Then, the systems were minimized, followed by two equilibration steps. In the first step, the temperature was equilibrated at 300 K over 1 ns of simulation under NVT ensemble. The Bussi thermostat was used with separate heat bath couplings for solute and solvent [26]. The second equilibration step was performed over 1 ns under NPT ensemble to keep the pressure at 1 bar employing the Parrinello-Rahman barostat [27].

We assessed the impact of sampling frequency and overall simulation length on the sampling of protein backbone conformations on five different proteins (PDB entries 1CLL, 1D3Z, 1QM3, 5PTI, and 2KUY, Supplementary Fig. 4). All simulations were performed for 1,000 ns and coordinates were recorded every 1 ns to limit the correlation between subsequent sets of coordinates. To assess whether the sampling time was sufficient to capture experimentally observed conformational states, we compared the conformational diversity in the MD trajectories against structures annotated in the CoDNaS database [28]. For the 22 proteins represented in the database, we found that the MD sampling either exceeds the conformational variability recorded in CoDNaS or produces similar results (Supplementary Fig. 5).

**Figure 4:**
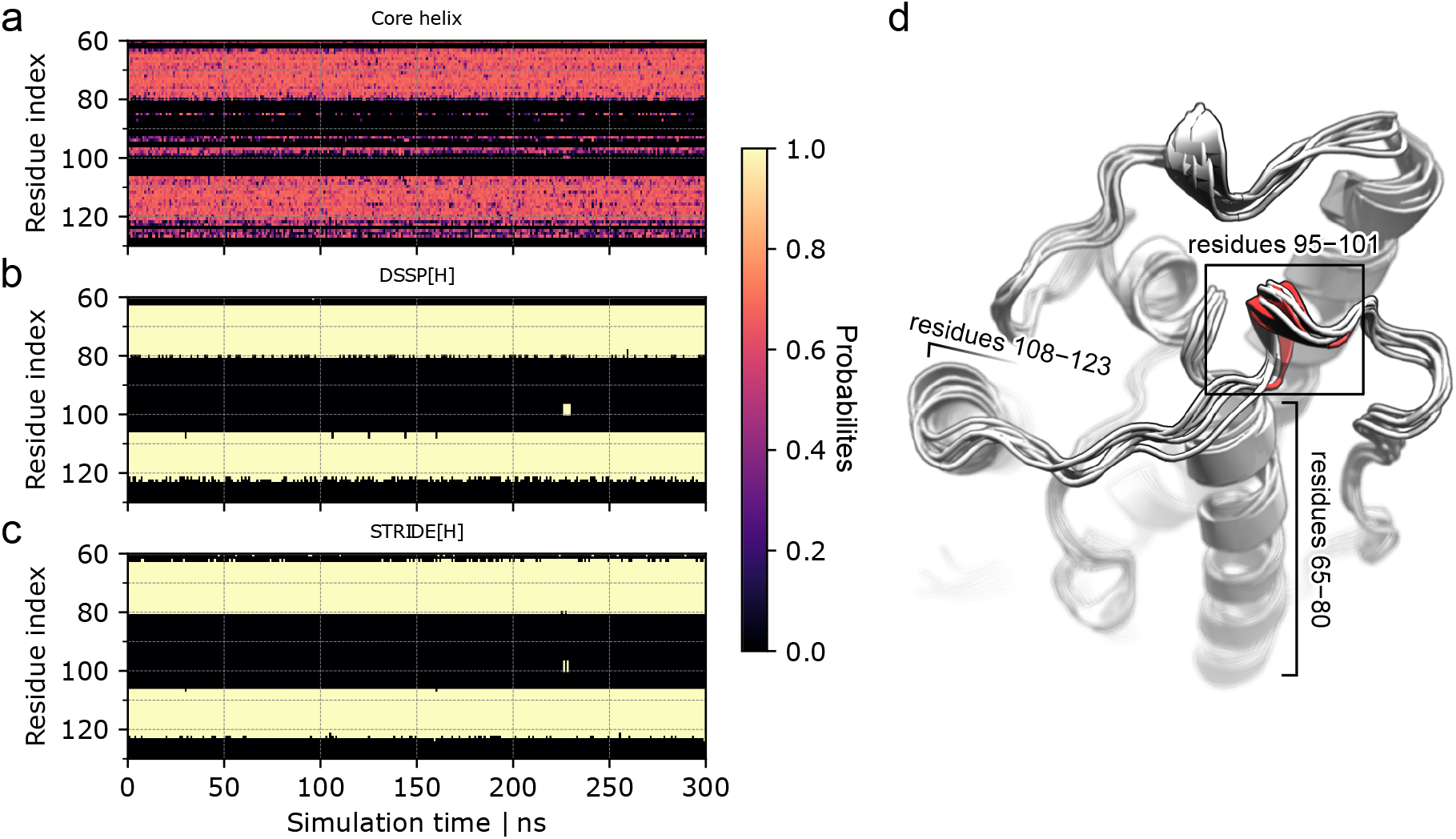
Comparison of conformational state propensities with traditional secondary structure assignments. The plot shows residues 60-130 of Endonuclease V (PDB ID: 2end). a) Assignments of *Core helix* propensities for the first 300 ns of the simulation. Notably, Constava con- tinuously detects *Core helix* propensities ∼ 0.6 for residues 95-100. b) DSSP assignment of H (α-helix) for the first 300 ns of the simulation. As DSSP performs a classification, the propensities for H are 0 or 1. The transient helix for residues 95-100 only appears shortly after more than 200 ns of simulation. c) STRIDE assignment of H (α-helix) for the first 300 ns of the simulation. As STRIDE performs a classification, the propensities for H are 0 or 1. The transient helix for residues 95-100 only appears shortly after more than 200 ns of simulation. d) Structure of Endonuclease V with regions shown in panels a, b, and c labeled.

**Figure 5:**
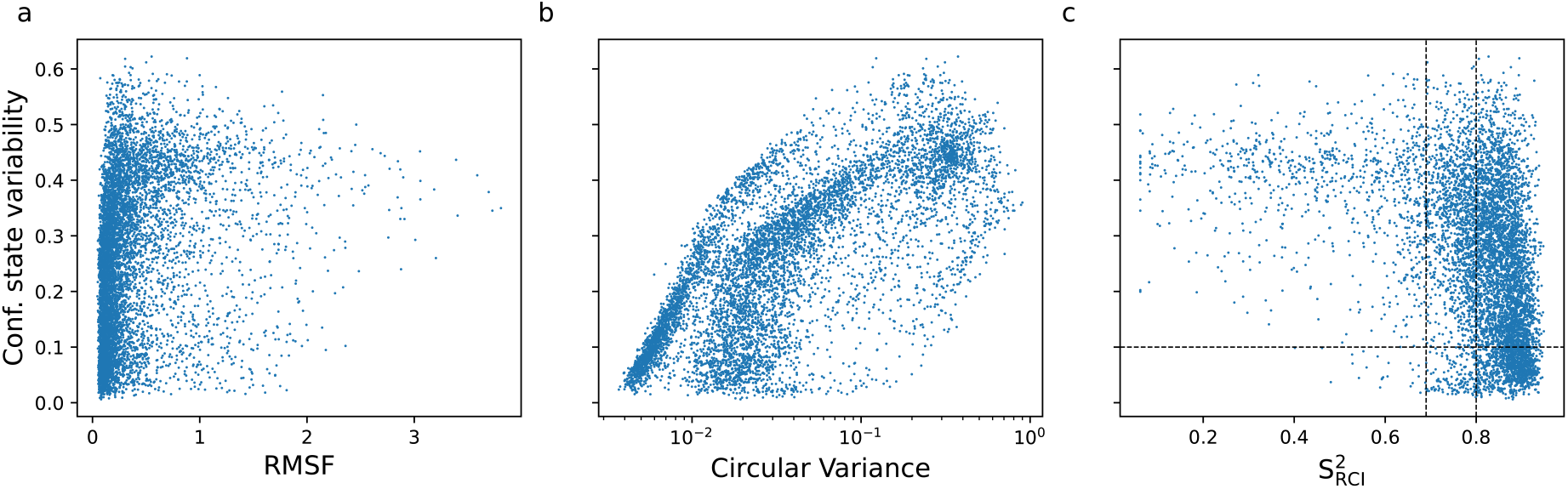
Conformational state variability (bootstrap sample size 3, 10,000 samples) versus traditional metrics. a) Root mean square fluctuations (RMSF) per residue calculated for all residues of proteins in the MD data set, Pearson’s *r* = 0.27 (p-value = 0.00). b) Circular variance (CV) calculated for all residues of proteins in the MD data set, Pearson’s *r* = 0.56 (*p <* 0.001). c) 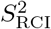 from residues in 62 proteins from the MD data set for which 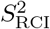 values were available (Supplementary Data 2), Pearson’s *r* = −0.41 (*p <* 0.001). The vertical dotted lines indicate the border between likely disordered, context-dependent and ordered residues (left to right) as defined in [17]. The horizontal dotted line is a visual guide to distinguish between residues with very low and higher conformational state variability.

To calculate the conformational state propensities and variability for each residue, the backbone (*ϕ, ψ*)-angles were extracted for all trajectories using GROMACS’ chi module. Then, conformational state propensities and variability were inferred as described above; samples of size 3 and 25 were drawn by bootstrapping for 10,000 times from the trajectory. Bootstrapping was chosen as with long sampling intervals independence of subsequent samples is assumed. Sampling size 3 was used to analyze local dynamics while sample size 25 informs on the conformational state propensity.

For comparison with traditional secondary structure assignments, we applied DSSP [29, 30] and STRIDE [19, 20] to all conformations extracted from the trajectories. For DSSP we used GROMACS’ do dssp module, while for STRIDE all frames of the MD trajectories were extracted as individual PDBs to be analyzed. We further calculated DSSP and STRIDE propensities as the fraction of frames per residue that are assigned to a given secondary structure category (H, E, …).

### Analysis of α- and β-synuclein

We obtained structural ensembles for both α-synuclein (αSyn) and β-synuclein (βSyn) from the Protein Ensemble Database (αSyn: PED00024, βSyn: PED00003) [31, 32, 33]. The backbone (*ϕ, ψ*)- angles were extracted from the conformational ensembles. To calculate the conformational state propensities and variability for each residue as described above, samples of size 25 were drawn by bootstrapping for 500 times.

## Results

In this section, we discuss application cases for our probabilistic approach to define protein conformation and dynamics. We will describe the relation between the metrics and established measures of conformation, and finally apply the method to structural ensembles derived from MD simulations and NMR experiments.

### Probabilistic definition of conformational states

The definition of the conformational states is based on the analysis of NMR ensembles, using the assumption that residues with a consistent structural definition in all models of the NMR ensemble adopt a single well-defined conformation, whereas residues with variable structure definitions adopt a wider range of conformations. This definition is further refined based on a ShiftCrypt-based interpretation of chemical shift information, which reflects the in-solution behavior of residues. In total 1,322 NMR structural ensembles (62,125 residues) were so analyzed and their residues subdivided into six conformational state categories (see Table 1): *Core helix, Surrounding helix, Core sheet, Surrounding sheet, Turn*, and *Other*. To obtain probabilistic models of these states in the backbone dihedral space, as extracted from NMR ensembles, Gaussian kernel density estimators (KDE) were trained. The conformational state propensities obtained with the dihedrals and each of the KDEs are a quantitative measure of the likelihood of a residue to assume this conformational state at any time of observation, given its backbone dihedral angles. These conformational states so reflect typical experimentally observed low energy states of residues (Fig. 2). The corresponding backbone dihedral space distribution indicates the expected range of backbone dihedrals that are available to a residue in that conformational state without having to overcome significant energy barriers. In particular the *Core helix* and *Core sheet* states exhibit deep wells in their energy landscapes, that in turn reflect the restrained backbone dynamics of such residues. In contrast, energy wells in *Turn* and *Other* states are more shallow, allowing for higher conformational freedom. A residue’s conformational state, thus, captures the preferential conformation of residues, as traditional secondary structure classification does, as well as dynamical properties of the local backbone.

**Table 1:** Description of conformational states and conformational state variability.

| Parameter | Abbreviation | Definition | Traditional interpretation |
| --- | --- | --- | --- |
| Conformational state variability | - | The change in conformational states probability vectors based on the dihedral angles of models in a protein structure ensemble. Residues with changing conformational states propensities within the different structures of the ensembles feature higher values. | The capability of a residue to adopt diverse conformational states. |
| *Core helix* | coreHelix | Residues with minimal backbone dynamics that exclusively exhibit α-helical conformation. | Residues in very stable rigid α-helices. |
| *Surrounding helix* | surrHelix | Residues with significant backbone dynamics that mostly exhibit α-helical conformation, but undergo conformational changes. | Residues found in kink regions of α-helices, but also encompassing transient helices, flexible helix termini, π-, and $3_{10}$-helices. |
| *Core sheet* | coreSheet | Residues with minimal backbone dynamics that exclusively exhibit β-sheet conformation. | Residues in stable rigid β-sheets but also in fully extended regions. |
| *Surrounding sheet* | surrSheet | Residues with significant backbone dynamics that mostly exhibit β-sheet conformation, but undergo conformational changes. | Residues found in kink regions and in the flexible termini of β-sheets as well as residues that preferably adopt the ppII state. |
| *Turn* | Turn | Residues with high backbone dynamics labeled as mostly Turn conformation by STRIDE. | Flexible residues found in turn-like conformations. |
| *Other* | Other | Residues with high backbone dynamics labeled as mostly Coil conformation by STRIDE. | Flexible residues that cannot be properly described by any of the other classes, typically found in disordered regions. |

We further define the conformational state variability parameter as the ability of a residue to adopt multiple conformational states in a given protein in a specific environment. This variability does not reflect actual dynamics, but rather indicates whether that residue is switching between different low energy conformational states. For example, a residue in states *Turn* or *Other* may exhibit high backbone dynamics (its (*ϕ, ψ*)-angles change constantly) while still having a low conformational state variability (its overall behavior is best described by one conformational state). This would be a typical conformational state adopted by residues in disordered regions of proteins. A residue in a well-defined stable helix, on the other hand, will also have low conformational state variability as well as very low backbone dynamics.

### Constava captures transient conformational states from MD ensembles

The new metric was tested on a diverse set of conformational ensembles extracted from MD simulations of 1,000 ns length for 113 proteins exhibiting varying degrees of structural order/disorder. From the trajectories (*ϕ, ψ*)-angles were extracted at intervals of 1 ns, to minimize the correlation between subsequent snapshots. The conformational state propensities were then inferred from each individual (*ϕ, ψ*)-angle pair, or using subsets of (*ϕ, ψ*)-angle pairs of varying sizes (Fig. 3a and b).

To investigate the impact of the number of (*ϕ, ψ*)-angle pairs in the inference, a bootstrapping procedure was first employed to sample *N* (*ϕ, ψ*)-angle pairs from random time points from the MD simulations. Increasing the number of *N*, where more angle pairs are considered at the same time, leads to residues being assigned more preferentially to specific conformational states. This is due to the conformational states sharing overlapping regions in the (*ϕ, ψ*)-space, so that a single angle pair data point will correspond to multiple regions (Fig. 2). When increasing the sample size *N*, however, the information available to the model to assign a specific conformational state increases and results in more unique assignments of the conformational state. Further increases of *N* also then start to bias unique conformational states, with lower populated states that the protein might transition through disregarded. Thus, to assess a protein’s conformational state variability lower values of *N* are more informative, whilst higher values reflect the most preferred conformational state(s). Based on our analysis of the impact of the sample size *N* on the results, we propose two sampling sizes also used throughout our work: *N* = 3 to study conformational state variability, as it provides the best compromise between a linear distribution values and a reasonable data range (Supplementary Figs. 2 & 3), while *N* = 25 is well suited to obtain preferred conformational states without losing information on less populated states (see Supplementary Fig. 6).

**Figure 6:**
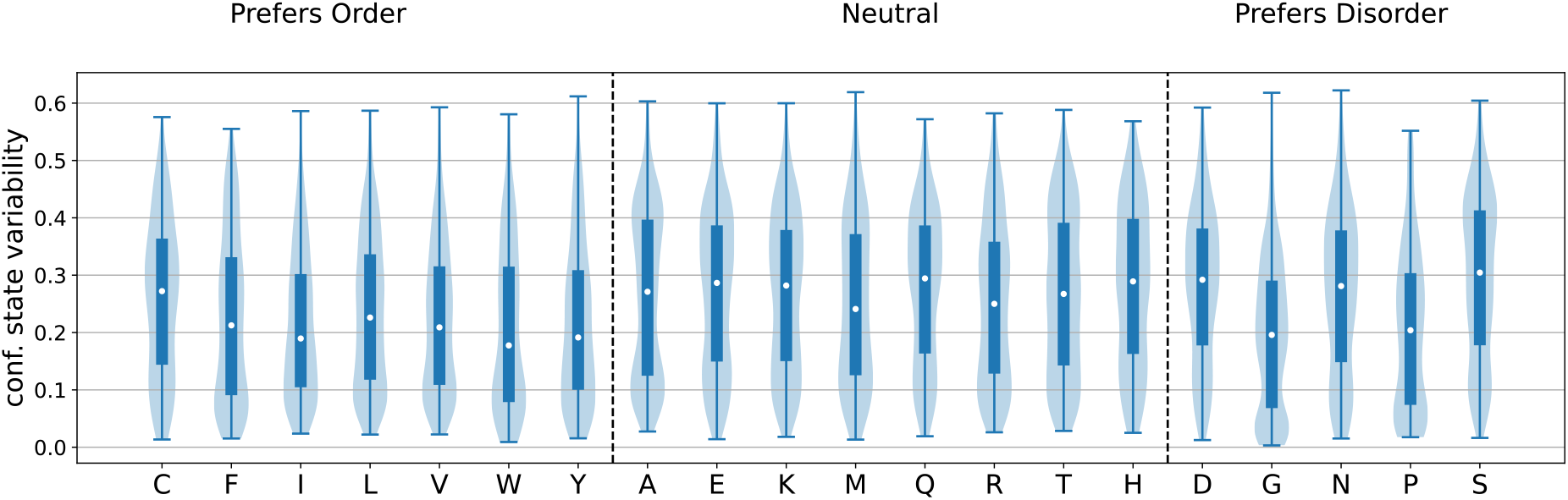
Conformational state variability values per amino acid type grouped according to their order preference. The 5 C- and N-terminal residues of each protein were excluded to remove end- of-chain bias towards disorder (*e*.*g*., methionine is often found as the first residue in a sequence). Amino acids which prefer order generally have the lowest conformational state variability. Glycine and proline have lower conformational state variability values as they tend to adopt respectively *Turn* and *Other* states. The raw data plot is available in Supplementary Fig. 11. The conformational state variability (Variability) displayed in this figure was inferred with bootstrap sample size 3 (10,000 samples). Each violin represents the density of residues along the range conformational state variability values. Encased within each violin, a blue bar delineates the inter-quartile range (IQR), extending from the first quartile (Q1) to the third quartile (Q3), thus, encompassing the middle 50% of the data points, and it contains a white dot which marks the median. From the ends of this bar, thin lines stretch out to the extremes, capped at the minimum and maximum values observed in the data set.

The random sampling of the bootstrapping procedure, however, loses any information about connections between models, such as the timeline in an MD simulation. A sliding-window approach was therefore implemented, which instead of randomly sampling from the whole population chooses blocks of adjacent (time-related) samples. This resulted in a much more sensitive detection of transient conformational states than regular bootstrapping (Fig. 3c and d). Moreover, by gradually extending the window size of the sliding window, information about the time intervals in which these states are transitioning between themselves can be obtained. Small window sizes pick up more transient states of shorter duration, while longer window sizes only detect those states that are stable for a longer time (Fig. 3c and e). Due to the higher sensitivity, using a sliding window is the preferred sub-sampling method for data obtained from time series (*e*.*g*., MD trajectories). For ensembles where no relation between adjacent ensemble members can be assumed (*e*.*g*., NMR ensembles), bootstrapping is the method of choice, as there is no direct connection between the sampled models.

### Relation between NMR-defined conformational states and traditional secondary structures assignments

DSSP [29, 30] and STRIDE [19, 20] are two of the most common tools to assign static secondary structure on protein structures. By applying them on every structure of an ensemble, conformational variable regions would receive changing assignments. To compare how such serial secondary structure assignments differ from our probabilistic method, we applied Constava, DSSP, and STRIDE to 113 MD ensembles. For the narrowly defined *Core helix* and *Core sheet* conformational states, near linear relations to H (α-helix) and E (β-sheet), respectively, emerged (Supplementary Fig. 7a,c & Supplementary Fig. 8a,c). This demonstrates that high propensities of *Core helix* and *Core sheet* indicate rigid residues that adopt stable secondary structure conformations, which are equally captured by DSSP and STRIDE. For the more dynamic conformational states, relationships with traditional secondary structures contributions are more complex, without clear correlations (Supplementary Fig. 7b,d-f & Supplementary Fig. 8b,d-f).

**Figure 7:**
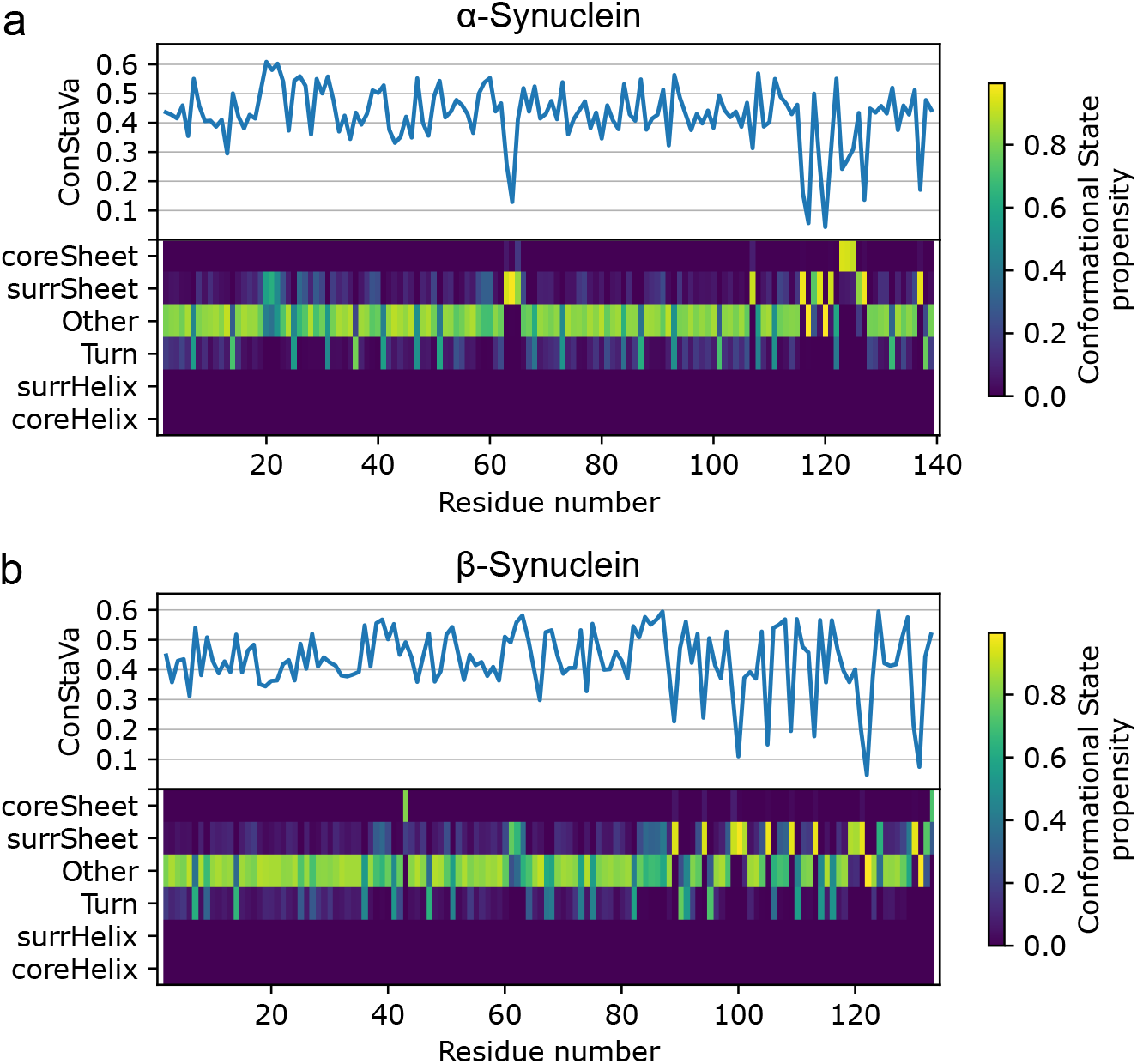
Results of Constava for α- and β-synuclein. For each protein the per-residue confor- mational state variability is shown (top) as well as the conformational state propensities for all the six conformational states (bottom) as calculated from the PED ensemble. a) In α-synuclein the N-terminal region (aa 1-98) is mostly in *Other* with intermittent *Turn* and localised *Surrounding sheet* conforma- tional states around residues 20 and 63. In the C-terminal part of the protein *Surrounding sheet* and *Core sheet* become more prominent, indicating an increased preference for extended structures, with reduced conformational state variability. b) In β-synuclein the N-terminal region (aa 1-76) is mostly in *Other* with again intermittent *Turn* conformational states and localised *Surrounding sheet* and one outlier *Core sheet* residue. The C-terminal part of the protein shows *Surrounding sheet* but no *Core sheet*, suggesting a prevalence of ppII-like conformations rather than actual β-sheets.

However, the probabilistic nature of the conformational states makes Constava more robust in detecting secondary structure propensities from ensemble data than approaches that only consider static protein structures like DSSP and STRIDE . When analyzing conformational ensembles from *e*.*g*. MD simulations, Constava has the potential to detect residues moving towards a secondary structure state, well before it is recognised by DSSP or STRIDE (Fig. 4 and Supplementary Fig. 9). In this example, Constava picks up on the propensities of multiple consecutive residues to form a helix long before the helix, as defined by DSSP and STRIDE, emerges. In the study of IDPs that exhibit high conformational freedom, so that relevant transient conformations cannot be efficiently sampled, this ability of Constava could be extremely useful in picking up pre-sampled conformations that are for example relevant for forming interactions, reducing the need for unachievable exhaustive sampling.

### Relation of conformational state variability to traditional dynamics metrics

The MD simulations enable a comparison of the conformational state variability with established metrics. The root mean square fluctuations (RMSF) and circular variance (CV), as calculated from the MD simulations, and the *S*^2^ order parameter derived from the NMR chemical shifts using the random coil index 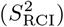 are shown in relation to the conformational state variability, calculated with the bootstrap method for 3 samples repeated 10,000 times, in Fig. 5 and Supplementary Fig. 10. There is a very weak correlation between RMSF and conformational state variability (see Fig. 5a), depicting a tendency for residues with low RMSF to also show low conformational state variability (*e*.*g*., in rigid stable helix). Whereas RMSF quantifies all backbone displacements, even if they do not affect the conformational state, the conformational state variability only measures switches between conformational states at the residue level.

There is a more pronounced correlation between CV and conformational state variability (Fig. 5b). CV represents the variation in the backbone dihedral angle space, with no incorporation of their conformational meaning. In contrast, conformational state variability represents transitions between conformational states that do not directly correlate with changes in the backbone dihedrals, as each conformational state accounts for varying degrees of backbone dynamics. Therefore, higher circular variance does not necessarily lead to high conformational state variability. For instance, a highly flexible coil would consistently occupy the *Other* state (low conformational state variablity), while displaying high backbone dynamics (high CV). For such a residue to show high conformational state variability as well, it would need to switch conformational states, e.g., by partaking in the formation of a transient helix.

The 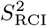, a measure of backbone rigidity derived from chemical shifts for the protein in solution, demonstrates a negative correlation to the conformational state variability (see Fig. 5c). Yet, the relationship is not directly linear. The cluster of residues with low conformational state variability and high 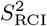 values (bottom right cell) represents residues with neither backbone movement nor changes in the conformational state space, such as in rigid stable helix. The region with 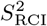 values below 0.69 predominantly features high conformational state variability values (top left cell), indicating highly dynamic residues that do change their conformational state. Note that there are also residues that have low conformational state variability, indicating they adopt a unique state (*Other* or *Turn*).

Finally, many residues have 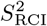 higher than 0.69, indicating residues where the NMR chemical shift values point to low backbone dynamics, but that still exhibit high conformational state variability. These represent residues that transition through the conformational state space but where these changes are not captured by the NMR chemical shift values (top middle and right cells). Since NMR chemical shift values are averaged over all conformations adopted by the protein in solution, this is possible when the chemical environment of the atoms in the residue remains relatively constant (*e*.*g*., switches between core and *Surrounding helix*) or where a rigid stable conformation dominates, but is not exclusive (*e*.*g*., a residue that is *Core sheet* in 50% of the conformations but adopts other conformational states the rest of the time).

These findings indicate that conformational state variability offers complementary insights that are not directly inferable from existing dynamics metrics.

### Conformational state variability for amino acids per order preference class

Amino acids have intrinsic properties that predispose them to facilitate the formation of ordered or disordered regions in proteins. Based on a previously used classification [17] the conformational state variability was examined within three classes: preferentially ordered (cysteine, phenylalanine, isoleucine, leucine, valine, tryptophan, and tyrosine), neutral (alanine, glutamic acid, lysine, methionine, glutamine, arginine, threonine, and histidine) and preferentially disordered (aspartic acid, glycine, asparagine, proline, and serine) residues. As expected, amino acids with preference for order exhibit lower conformational state variability than those with preference for disorder (Fig. 6). Notable exceptions in the disordered category are glycine and proline. Both show low conformational state variabilities, which in this case reflects their tendency to exclusively adopt *Turn* or *Other* conformational states, without switching to other conformational states. Both these states represent mostly flat energy landscapes with low energy barriers, thus allowing for random movement or disorder (Fig. 2). Note that aspartic acid, asparagine and serine have higher conformational state variability, indicating they are more capable of switching states. This highlights an interesting feature of the conformational state variability, residues that remain in the same low energy conformational state, even if that conformational state is effectively dynamic, exhibit low conformational state variability. This is also reflected in *Other* conformational states in Supplementary Fig. 13.

### Conformational states capture regions of partial order in α- and β- synuclein

To assess the ability of our method to obtain relevant features from structural ensembles, we applied the method to two the Protein Ensemble Database (PED) ensembles of two well studied IDPs: α-synuclein (αSyn) and β-synuclein (βSyn). αSyn is an intrinsically disordered peptide whose aggregation plays a central role in the pathogenesis of Parkinson’s disease, while its βSyn family member is less prone to aggregation.

Both proteins have been extensively studied by NMR, which indicated that the N-terminal part of both proteins (αSyn: aa 1-98, βSyn: aa 1-76) adopts random coil conformation. In contrast, the C-terminal region of both proteins exhibited extended conformations, with αSyn primarily being in β-sheet conformation [32] and βSyn adopting a polyproline II (ppII) state [34]. Constava here illustrates its ability to extract these key features from a structural ensemble. While the N-terminal parts in both proteins primarily show *Other* with intermittent *Turn* corresponding to random coil state, the C-terminal propensities are shifted towards extended conformations, *i*.*e*., *Surrounding sheet* and *Core sheet* (Fig. 7). Notably, the *Core sheet* region, which is exclusively populated in αSyn is strictly limited to β-sheet conformations, while *Surrounding sheet* which is prevalent in βSyn covers all extended states including the ppII region. Whilst these ensembles were generated using the NMR data and so should reflect experimental observations, Constava here shows the ability to directly pick up these relevant structural determinants with straightforward visualisation and interpretation possible.

## Discussion

The advent of AlphaFold2 has revolutionised the field of structural bioinformatics by rapidly and accurately providing the structure of protein regions that adopt a stable well-defined fold. The focus is now shifting to predicting multiple conformational states for proteins, with as extreme case providing structure ensembles for intrinsically disordered regions or proteins. However, describing such conformational ensembles requires probabilistic metrics that capture (transient) features of such ensembles, as single models cannot represent the overall behavior of a protein, for example even if the protein does form transient helix turns these will traditionally only be identified in the models if stable hydrogen bonds are present. This work introduces a probabilistic data-driven definition of likely low-energy conformational states of amino acid residues in proteins based on the analysis of NMR ensembles and associated chemical shifts. We defined a total of six conformational states: *Core helix, Surrounding helix, Core sheet, Surrounding sheet, Other*, and *Turn*. We further introduced the concept of conformational state variability, a parameter that describes how often a residue switches between these conformational states. Both concepts are encompassed in the Constava software, which enables users to infer conformational states and their variability from an ensemble of structure models produced by molecular dynamics or other approaches. Constava describes the observed behaviour of each individual residue in such a conformational ensemble in terms of the relative propensities of the conformational states it adopts. Previous work demonstrated how such probabilistic descriptions can be used to efficiently sample realistic protein conformations [35] .

When applied to molecular dynamics simulations, Constava is able to capture preferred and transient conformational states adopted by residues. Small sample sizes detect low populated transient states and provide a description of the conformational state dynamics of the residue. In contrast, larger sample sizes will detect only preferential conformational states and disregard transient states. The method can therefore detect different conformational features depending on the parameters used. When the subsampling directly uses a sliding window over the time series information in MD trajectories, it is even more sensitive towards transient conformational states that residues adopt. Since the sample size in that case directly relates to time intervals in the simulation, the method can be fine tuned to capture conformational states that persist specific time intervals. This distinguishes Constava from predictors of disordered regions or aggregation regions, such as IUPred [36], DisoMine [37], or TANGO [38]. It provides values representing the likelihood for the defined conformational states, which also capture the protein’s conformational dynamics, while through the conformational state variability it identifies the way residues change between conformational states. Moreover, as these values are derived from statistical potentials, the Constava conformational states in principle allow for a thermodynamical interpretation (Fig. 2).

The conformational state variability definition is non-linear and is not necessarily directly related to absolute changes in backbone dihedral angle space. The *Other* state, for example, covers a wide range of dihedral space, so if a residue samples this range in dihedral space it will be exclusively assigned the *Other* state and have low conformational state variability - it adopts the same low-energy conformational state. Further motions within this same conformational state then also do not lead to any changes in its conformational state. Residues that exhibit high conformational state variability on the other hand do switch between different typical low-energy states. They therefore indicate regions that experience “conformational state switches” in the protein. Such an interpretation differs significantly from metrics based purely on structural descriptors like root mean square fluctuations (RMSF) or circular variance (CV), which non-discriminatively interpret variations in respectively inter-atomic cartesian distances or dihedral angles. Consequently, it is not possible to infer the conformational state variability directly from these other metrics (Fig. 5).

The conformational state variability does relate in different ways to the NMR chemical shift derived RCI-derived *S*^2^ order parameter 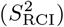. Three categories of residues can be discerned: Highly ordered residues with low potential for conformational states changes, disordered residues which continuously change between conformational states, and residues capable of adopting stable conformational states, but with the capability of transitioning among different states. The entropy of the conformational states in the (*ϕ, ψ*)-space is a measure of how sensitive the conformational state variability is to changes in the backbone dihedrals (Supplementary Fig. 14). Remarkably, the last category is comprised of residues with relatively high 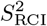, indicating little backbone dynamics, that still transition between states. This showcases how Constava can detect conformational state changes that are not picked up by NMR chemical shift values.

The conformational state variability also provides an interesting perspective on amino acid behavior. Although the general categories of order-neutral-disorder inducing amino acids are as expected represented by low to high conformational variability, glycine and proline frequently feature lower values than the other amino acids in the disordered category. This indicates their preference to adopt exclusively the *Turn* or *Other* conformations as their low-energy states, with wide and flat energy landscapes for their dihedral space that allow them to continuously adopt a wide range of backbone conformations (Fig. 2). Only rarely do they adopt multiple conformational states, in contrast to serine for example, which easily switches between them and can therefore accommodate multiple conformational states. Constava can, in this light, also identify regions with higher degrees of order in structure ensembles of intrinsically disordered proteins, represented by core and surrounding states. In the PED ensembles of α-synuclein (αSyn) and β-synuclein (βSyn), Constava directly identifies the β-sheet prone regions in αSyn as well as the ppII prone regions in the C-terminus of βSyn.

These examples showcase how the probabilistic definitions of conformational states allows for the reliable extraction of high-level information about protein behavior in the study of (highly) dynamic proteins. Where large conformational ensembles are available, Constava does offer a novel way to identify the conformational states of residues, as well as how likely residues are to switch between conformational states. This is particularly useful in describing and classifying proteins with highly variable structures, such as IDPs [1] or fold-switching proteins [39], for which static definitions of conformation are not appropriate. We hope this approach will help in redefining our interpretation of proteins as dynamic, not static, entities with variable behavior. The method is available as a PyPI package and all data used in this is freely available in the public domain.

## Data availability

All data used in this work have been compiled in a single Zenodo repository (https://doi.org/10.5281/zenodo.10371447).

## Code availability

The Constava software is available as a PyPI package (https://pypi.org/project/constava/). The source code is accessible on GitHub (https://github.com/Bio2Byte/constava) and may be used and redistributed under GPLv3 license; version 1.0.0 in preparation of this manuscript has been deposited in Zenodo (https://doi.org/10.5281/zenodo.10649793). An interactive Colab notebook is available with examples of the execution of the software https://colab.research.google.com/github/Bio2Byte/public_notebooks/blob/main/constava_examples.ipynb.

## Acknowledgements

We thank Adrian Diaz for the invaluable help in the distribution of this software. This work has been supported by the European Union’s Horizon 2020 research and innovation program under the Marie Sklodowska-Curie grant agreement [813239 to J.R.-M. and J.G.-G.]; Research Foundation Flanders (FWO) [G.032816N to G.O., G.028821N to D.B.]; Research Foundation Flanders (FWO) International Research Infrastructure [I000323N to W.V.]; COST Action ML4NGP, CA21160, supported by COST (European Cooperation in Science and Technology). The resources and services used in this work were provided by the VSC (Flemish Supercomputer Center), funded by the Research Foundation - Flanders (FWO) and the Flemish Government.

## Author contributions

W.V., D.R. and G.O. conceptualised the study. G.O. developed the initial methodology. W.V., D.R., G.O. and D.B. provided supervision. W.V. provided the NMR data and directed the project. J.G.-G. and D.B. developed, implemented and validated the method. J.R.-M. performed the MD simulations. All authors contributed to the writing of the manuscript.

## Competing interests

The authors declare no competing interests.

## Notes

https://pypi.org/project/constava/

https://doi.org/10.5281/zenodo.10371447

### Competing Interest Statement

The authors have declared no competing interest.

### Summary of Updates

Figure 4 has been modified to expand the analysis and also STRIDE secondary structure assignments, and supplementary figure 9 was expanded to also include the STRIDE assignments. Materials and Methods was modified to reflect these changes. Supplementary figure 8 was also added as a STRIDE analogous figure to supplementary figure 7. Supplementary figure 15 added to indicate co-occurrence of conformational states.

