## Supplementary information for "Data-driven probabilistic definition of the low energy conformational states of protein residues"

This work is shared under a CC BY 4.0 license.

List of proteins mapped to NMR metrics

This work is shared under a CC BY 4.0 license.

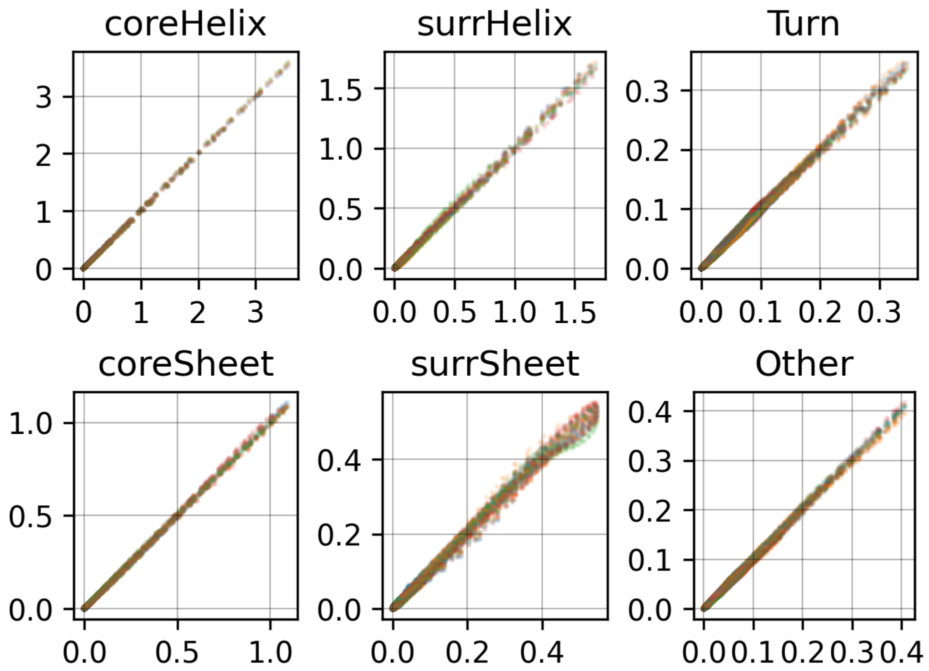

***Supplementary Fig. 1. Cross-validation of probability density kernels of conformational states.*** *The kernels were fitted five times, leaving out 20% of the ensembles each time. The plots show the correlation between probability densities sampled from the five different kernels.*

This work is shared under a CC BY 4.0 license.

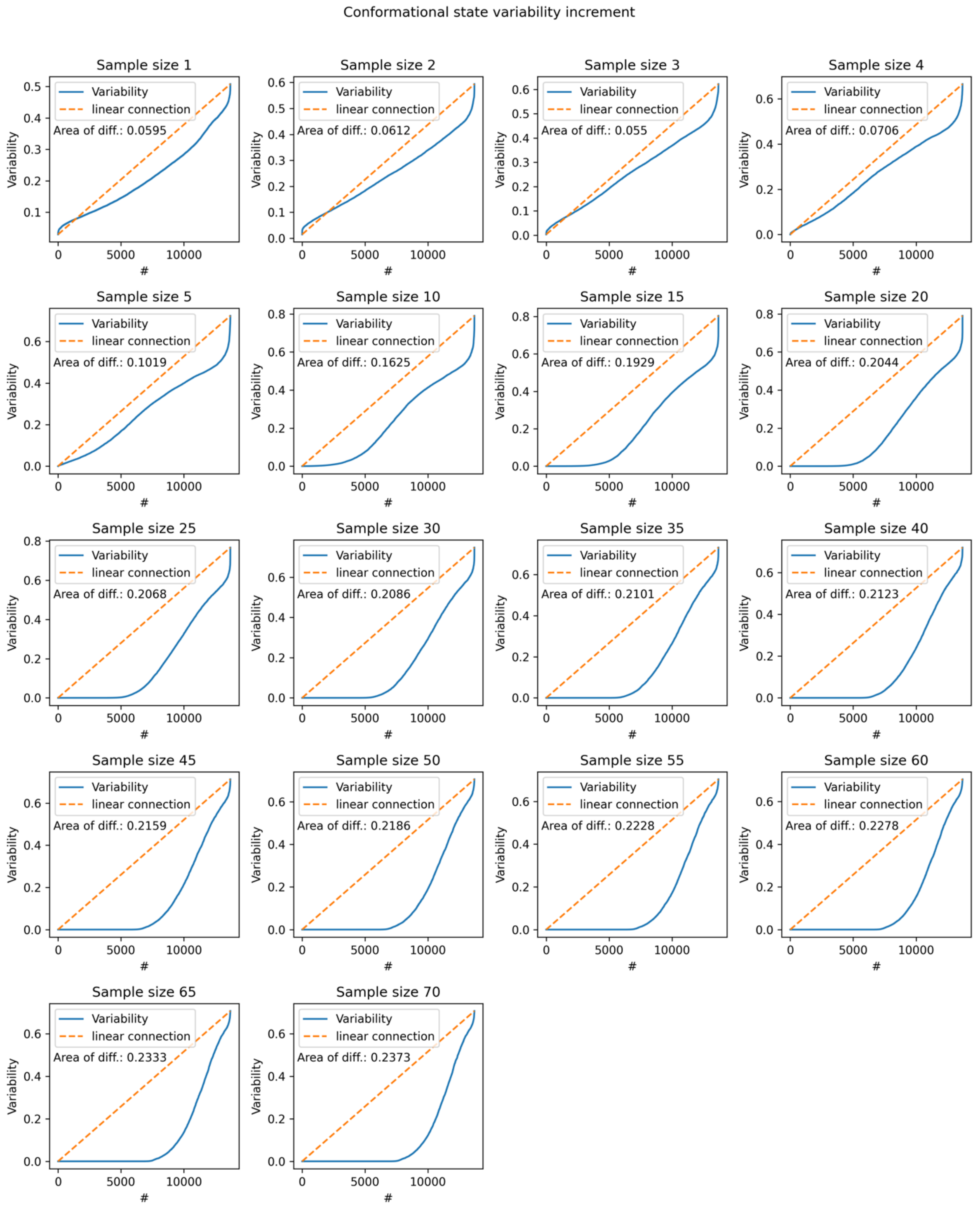

***Supplementary Fig. 2. Influence of the sample size on the conformational state variability.*** *Conformational state variabilities were calculated for all residues in the MD ensembles employing bootstrapping with different sample sizes (10,000 samples). The resulting values were plotted in ascending order (blue), while the dashed line (orange) represents an ideal line between the minimum and maximum values. Thus, the area between the two lines is a measure of the linearity of the conformational state variability in the dataset. As the sample size increases, less samples fall in the linear section of the conformational state variability metric. A sample size of 3 was chosen to obtain informative conformational state variabilities from ensembles for presenting the highest linearity in the metric while also spanning over a decent value range (min-max, see Supplementary Fig. 3).*

This work is shared under a CC BY 4.0 license.

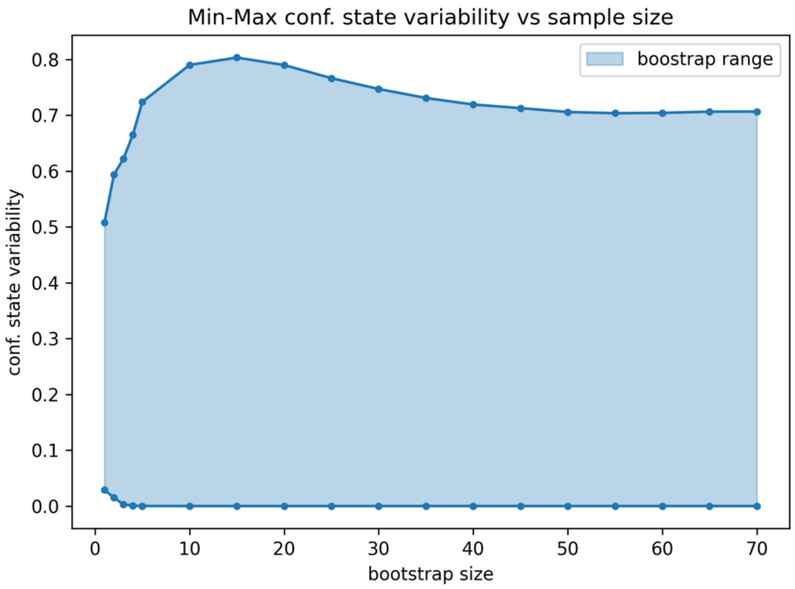

***Supplementary Fig. 3. Range of conformational state variability values as a function of the sample size.*** *Conformational state variabilities were calculated for all residues in the MD ensembles employing bootstrapping with different sample sizes (10,000 samples). The minimum and maximum values are shown here. Initially, as the sample size increases, the range between minimum and maximum values increases as well, reaching a maximum at a sample size of 15. For low sample sizes, conformational state variabilities of 0 are practically not observed. This indicates that even smallest fluctuations of the backbone dihedrals lead to an increase in conformational state variability. As the sample size increase, those small fluctuations become less important, allowing for conformational state variabilities that are practically 0.*

This work is shared under a CC BY 4.0 license.

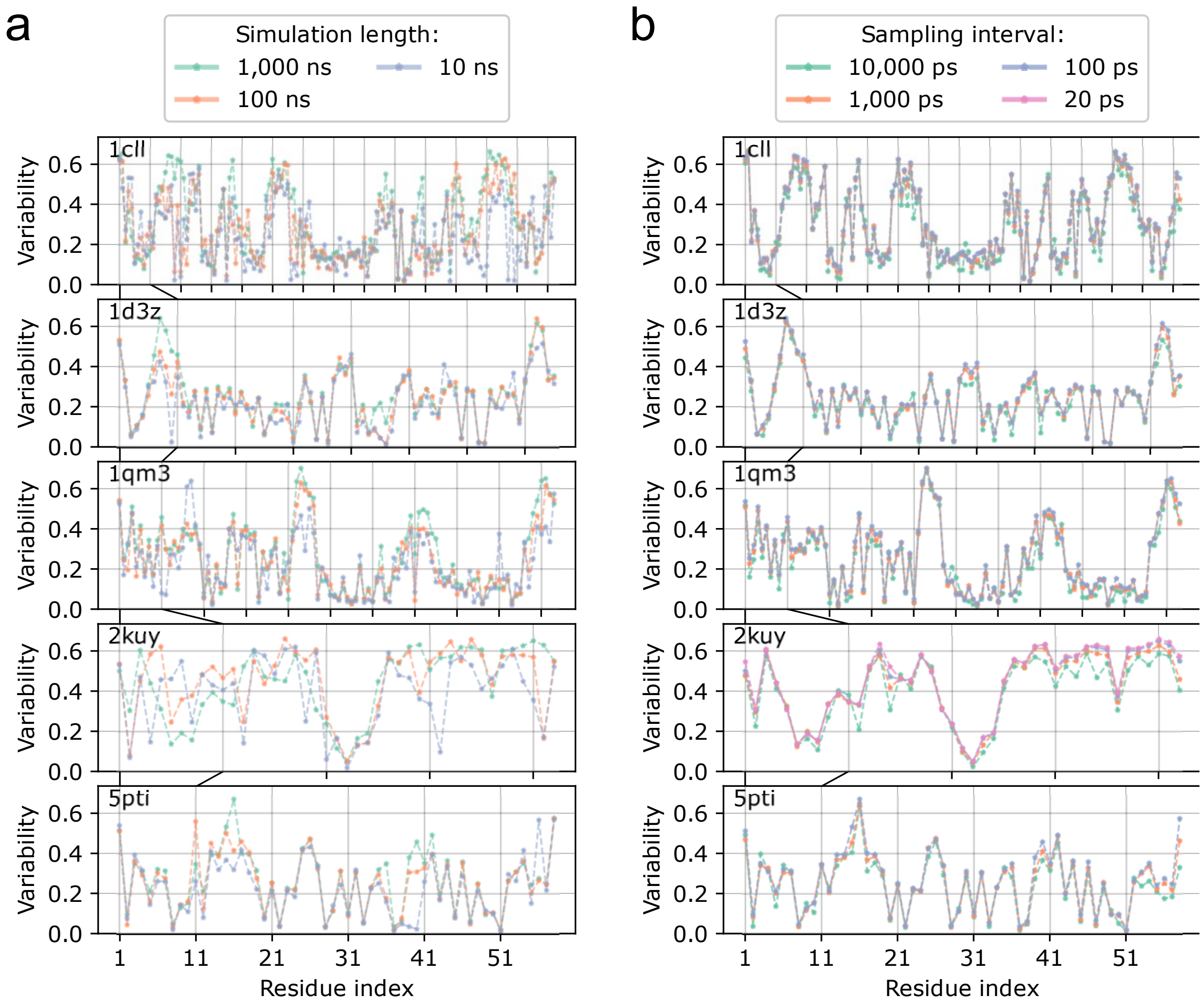

***Supplementary Fig. 4. Analysis of sampling parameters for molecular dynamics simulations.*** *Five proteins were simulated to assess the impact of different sampling parameters on the results. From all simulations the conformational state variabilities were calculated using the sliding window-approach with a window size of 3. Results are shown as the conformational state variability along the sequence for different sampling parameters. a) Analysis of the of the impact of simulation time. Results after 100 ns did not differ much from longer simulations. b) Analysis of the impact of sampling intervals. Only minor changes were observed. Sampling in intervals of 1,000 ps was chosen for the final simulations in this work, to minimize the autocorrelation between subsequent sampling time points.*

This work is shared under a CC BY 4.0 license.

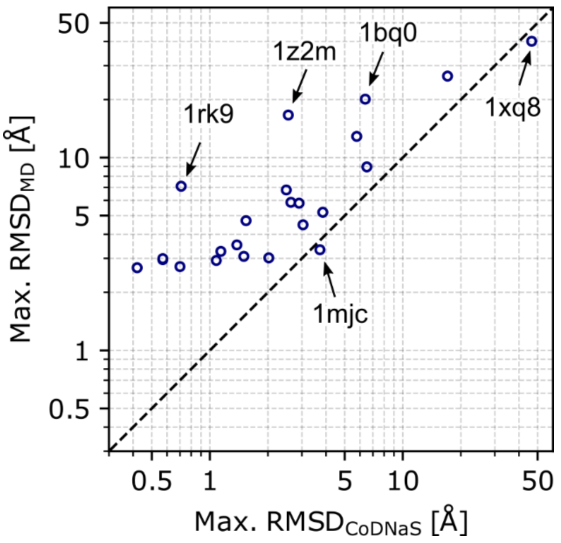

***Supplementary Fig. 5. Comparison of conformational diversity between MD ensembles and CoDNaS.*** *For a given protein the maximum RMSD values between any structures in CoDNaS (x-axis) and the MD ensemble (y-axis) are depicted. With two exceptions (1mjc and 1xq8) the MD simulations produced more diverse ensembles than the structures in CoDNaS. For some structures* (e.g., *1z2m and 1bq0) the conformational variability in the simulation strongly exceeded that in CoDNaS. This is mostly due to flexible termini, which MD simulations sample freely. A notable exception is 1rk9, which remains fully folded throughout the simulation but undergoes a conformational change which results in a higher RMSD than observed in CoDNaS. Out of the 113 simulated proteins, only 22 are annotated in CoDNaS.*

This work is shared under a CC BY 4.0 license.

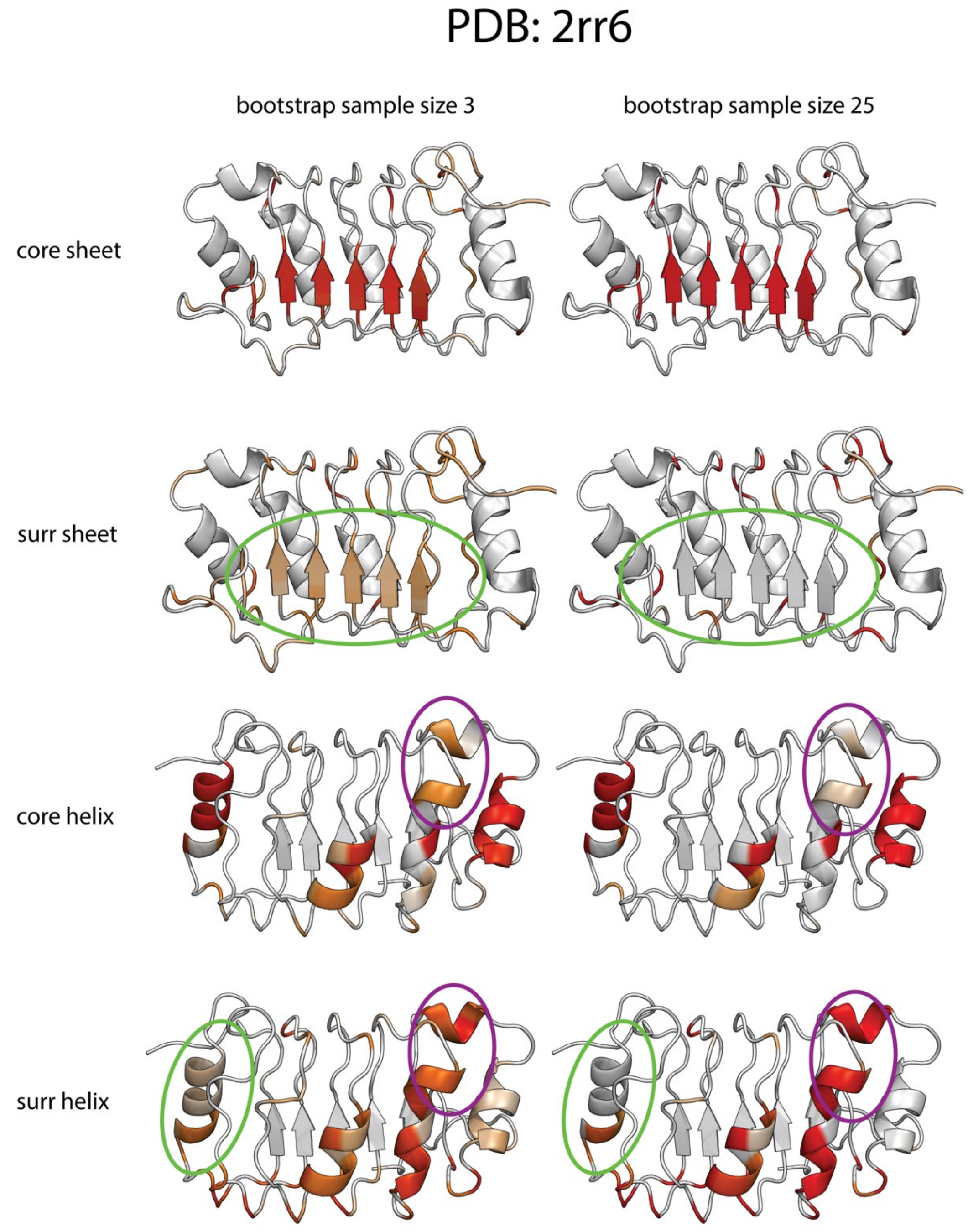

***Supplementary Fig. 6. Impact of the sample size on conformational states probabilities in human acidic leucine-rich nuclear phosphoprotein 32 family member B (PDB ID: 2rr6).*** *Conformational state propensities were inferred from the MD ensembles employing bootstrapping with sample sizes 3 and 25 (10,000 samples). As the sample size increases, ambiguous assignments (shades of orange) become less apparent, rather a preferred conformational state is assigned with high propensity (shades of red). On the first two rows, the low probability for Surrounding sheet in the circled area disappears in favour of higher probabilities towards Core sheet. Similarly, in the last two rows the green circled region experiences an increase in Core helix probability. Contrarily, the purple circled region experiences an increase Surrounding helix probability in detriment of Core helix probability.*

This work is shared under a CC BY 4.0 license.

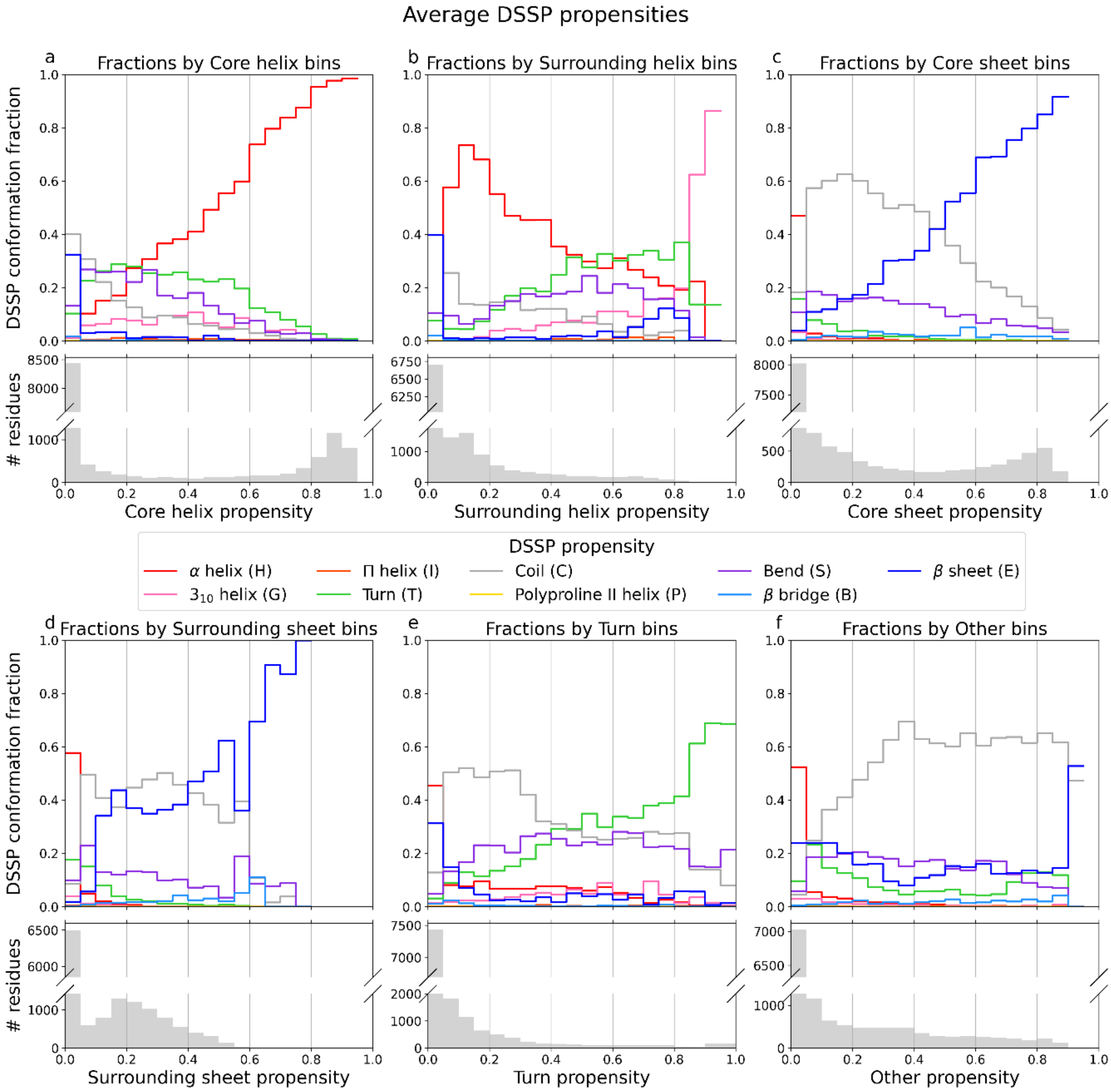

***Supplementary Fig. 7. Relation between Constava conformational states and DSSP secondary structure assignments.*** *Constava conformational state propensities were correlated with DSSP propensities (average fraction of frames assigned to a given secondary structure by DSSP). The two classification schemes were compared in bins. The bar plot below the correlation plots shows the number of samples (residues) in each bin. a) For* Core helix *propensities there is a correlation to H (α-helix). b) For* Surrounding helix *propensities no clear correlations to any DSSP category can be observed. The values for bins with propensities > 0.8 cannot be properly interpreted due to the low number of samples. c) For* Core sheet *there is a correlation with E (β-sheet). d) For* Surrounding sheet *no clear correlations are apparent. There is some correlation with E, but the relationship is non-linear and the low number of samples for propensities > 0.6 makes the interpretation difficult. e) For* Turn *a correlation with T (hydrogen bonded turns) emerges but other DSSP categories are observed for all levels of* Turn *propensity. f) Finally, the* Other *propensity mostly associates to C (coil), but also other DSSP categories are observed for all levels of* Other *propensity. Since* Turn *and* Other *indicate highly dynamic residues, it makes sense, that these residues form other secondary structure elements transiently.*

This work is shared under a CC BY 4.0 license.

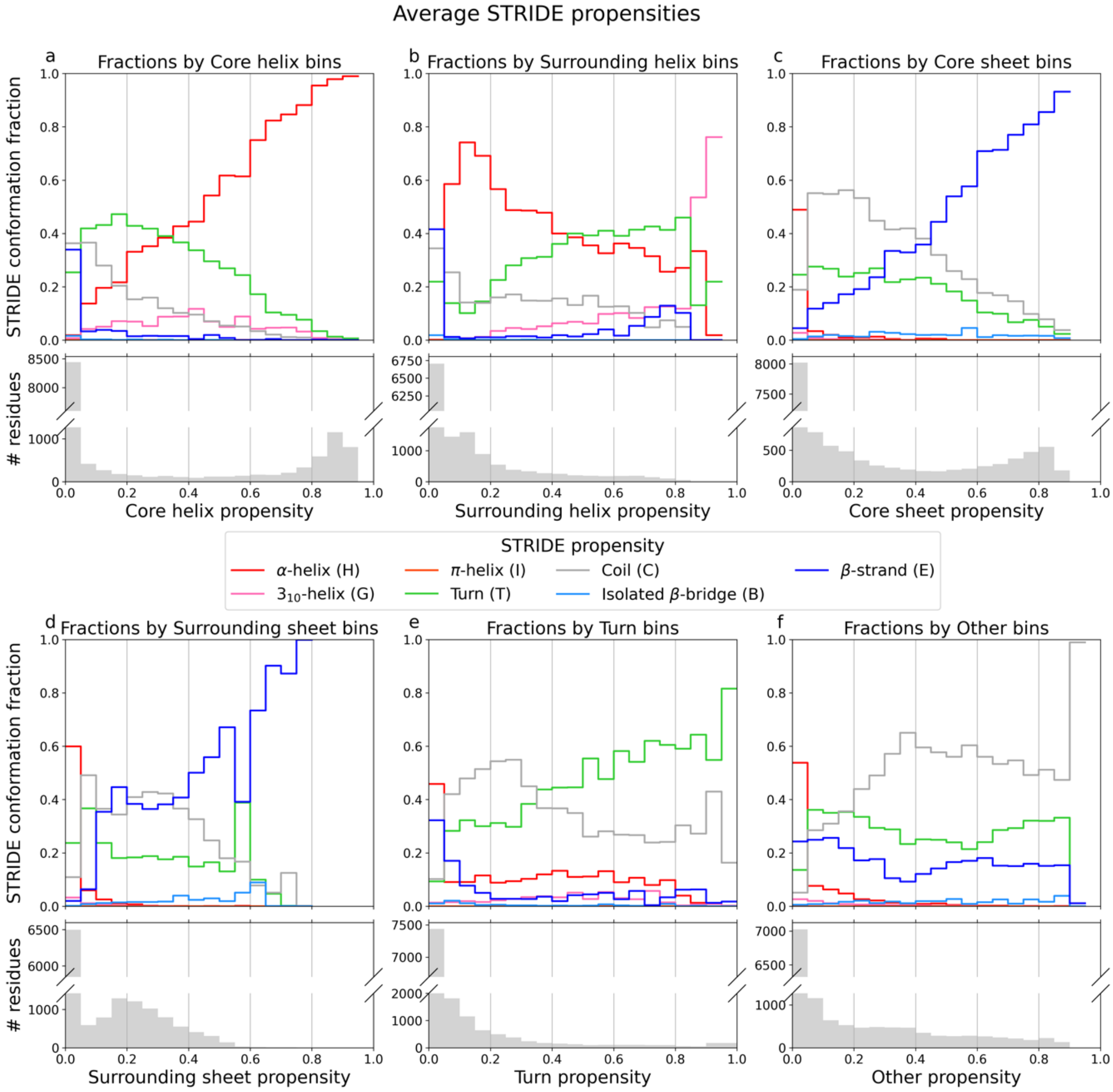

***Supplementary Fig. 8. Relation between Constava conformational states and STRIDE secondary structure assignments.*** *Constava conformational state propensities were correlated with STRIDE propensities (average fraction of frames assigned to a given secondary structure by STRIDE). The two classification schemes were compared in bins. The bar plot below the correlation plots shows the number of samples (residues) in each bin. a) For* Core helix *propensities there is a correlation to H (α-helix). b) For* Surrounding helix *propensities no clear correlations to any STRIDE category can be observed. The values for bins with propensities > 0.8 cannot be properly interpreted due to the low number of samples. c) For* Core sheet *there is a correlation with E (β-sheet). d) For* Surrounding sheet *no clear correlations are apparent. There is some correlation with E, but the relationship is non-linear and the low number of samples for propensities > 0.6 makes the interpretation difficult. e) For* Turn *a correlation with T (hydrogen bonded turns) emerges but other STRIDE categories are observed for all levels of* Turn *propensity. f) Finally, the* Other *propensity mostly associates to C (coil), but also other STRIDE categories are observed for all levels of* Other *propensity. Since* Turn *and* Other *indicate highly dynamic residues, it makes sense, that these residues form other secondary structure elements transiently.*

This work is shared under a CC BY 4.0 license.

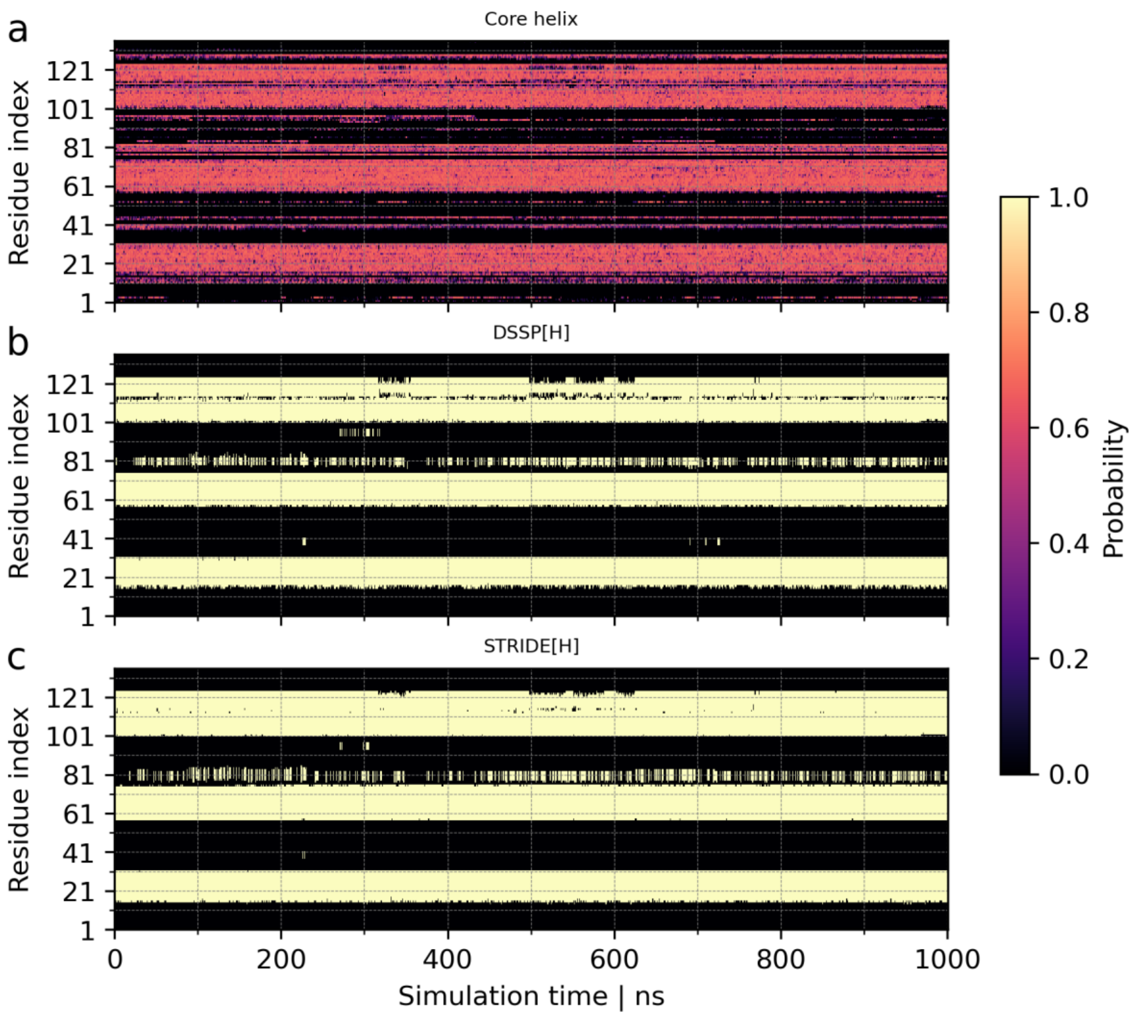

***Supplementary Fig. 9. Comparison of Constava and traditional secondary structure assignments with respect to their ability to detect transiently forming secondary structure elements.*** *Constava, DSSP, and STRIDE were applied to all conformations in the MD trajectories. Shown are: a) Core helix propensities, b) DSSP H (α-helix) assignments, and c) STRIDE H (α-helix) assignments along the MD trajectory of Endonuclease V (PDB ID: 2end). Beyond the relation between propensities and fractions in Supplementary Fig. 7 and Supplementary Fig. 8, Constava shows the capability to detect conformational states before the criteria for the other methods are met. In addition to returning high Core helix probability to the stable helix regions that DSSP and STRIDE assign, some regions with more dynamic behaviour interestingly show detection of helix propensity. The region between amino acids 95-101 is detected to have notable Core helix propensity (a) by Constava throughout the totality of the simulation, even though it is rarely assigned by DSSP (b) and STRIDE (c) in the MD time steps. Additionally, the region between amino acids 39-45 shows a similar tendency, with a longer amino acid range of high Core helix propensities near in the 300 ns time step, where DSSP and STRIDE detect α-helix. This longer stretch indicates that enough amino acids to form the necessary hydrogen bonds for DSSP and STRIDE assignment have adopted Core helix conformation. Finally, the region between amino acids 55-60 is also detected along the whole simulation, while DSSP and STRIDE intermittently assigns α-helix.*

This work is shared under a CC BY 4.0 license.

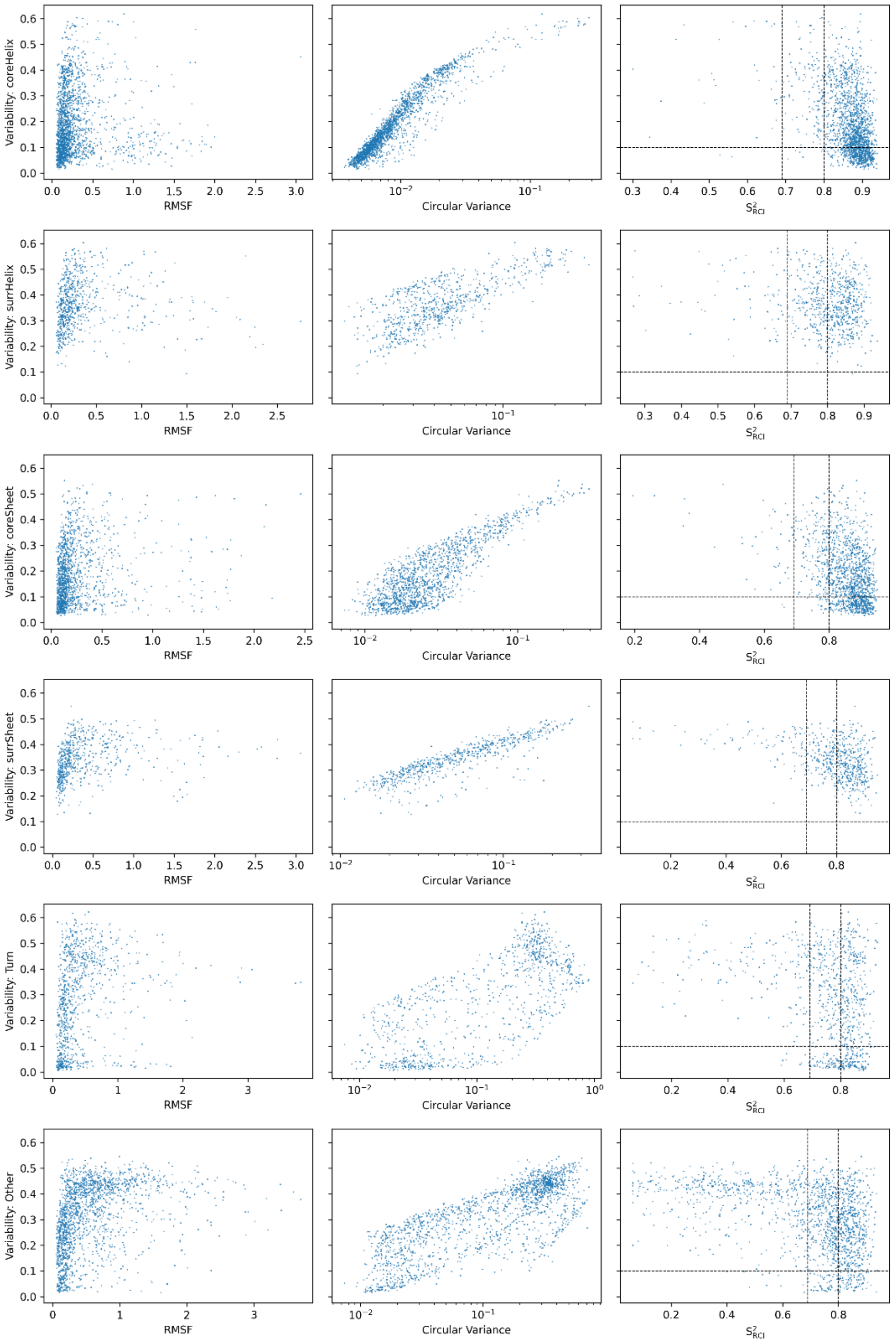

***Supplementary Fig. 10. Comparison of conformational state variability with traditional dynamics metrics per prevalent conformational state.*** *Conformational state propensities were inferred from the MD ensembles employing bootstrapping with sample sizes 3 (10,000 samples). The results were subdivided by their prevalent conformational state. From top to bottom these are: Core helix, Surrounding helix, Core sheet, Surrounding sheet, Turn and Other. In the left column, the conformational state variability is potted against the root mean square fluctuations (RMSF). In the central column, the conformational state variability is plotted against the circular variance. On the right, the conformational state variability is plotted against the $S^2_{RCI}$. The vertical dotted lines indicate the border between likely disordered, context-dependent and ordered residues (left to right) as defined in previous work [1]. The horizontal dotted line is a visual guide to distinguish between residues with very low and higher conformational state variability.*

This work is shared under a CC BY 4.0 license.

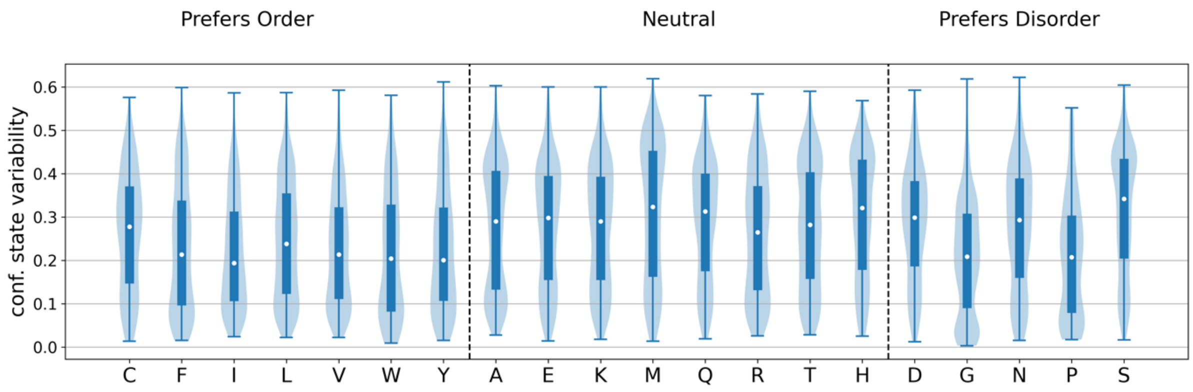

***Supplementary Fig. 11. Conformational state variability values per amino acid type according to their order preference, terminal regions included.*** *Conformational state variabilities were calculated for all residues in the MD ensembles employing bootstrapping with sample size 3 (10,000 samples). Amino acids were classified depending on their intrinsic order preference according to the classification on previous work[1] as preferentially ordered (cysteine, phenylalanine, isoleucine, leucine, valine, tryptophan & tyrosine), neutral (alanine, glutamic acid, lysine, methionine, glutamine, arginine, threonine & histidine) and preferentially disordered (aspartic acid, glycine, asparagine, proline & serine). Amino acids which prefer order generally featured lower conformational state variability values than the remaining. The homologous figure with terminal residues removed can be found in the main text, Figure 5. This figure mainly differs in the variability values distribution for methionine, which features higher values in this figure than in Figure 6. This is due to the preferential location of Methionine at the N-terminal region of the protein, which is less bound to inter-residue interactions and allows for easier conformational state changes and therefore variability.*

[1] Cilia, E., Pancsa, R., Tompa, P., Lenaerts, T. & Vranken, W. F. From protein sequence to dynamics and disorder with DynaMine. *Nature Communications* 4, 2741 (2013).

This work is shared under a CC BY 4.0 license.

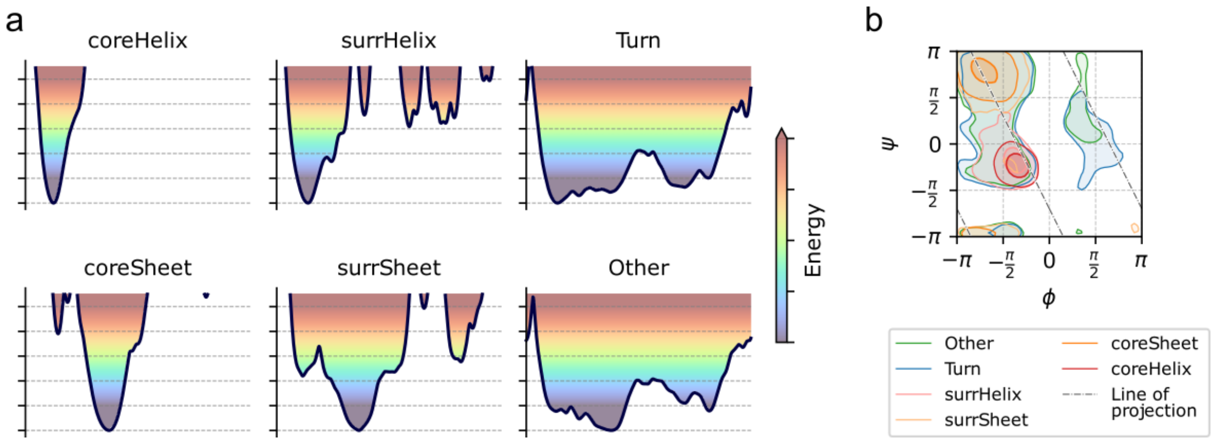

***Supplementary Fig. 12. Schematic representation of the conformational states as energy landscapes.*** *a) Cross section through the energy landscapes along the line of projection (shown in panel b). coreHelix and coreSheet conformational states are defined by a single deep energy well, vastly restricting backbone mobility. The states surrHelix and surrSheet show lowered energy barriers, allowing for more backbone mobility. Finally, in the Other and Turn conformational states, most of the dihedral space is accessible with only minor energy barriers in between. b) Schematic overview of the definition of the conformational states, as well as the line of projection used in panel a.*

This work is shared under a CC BY 4.0 license.

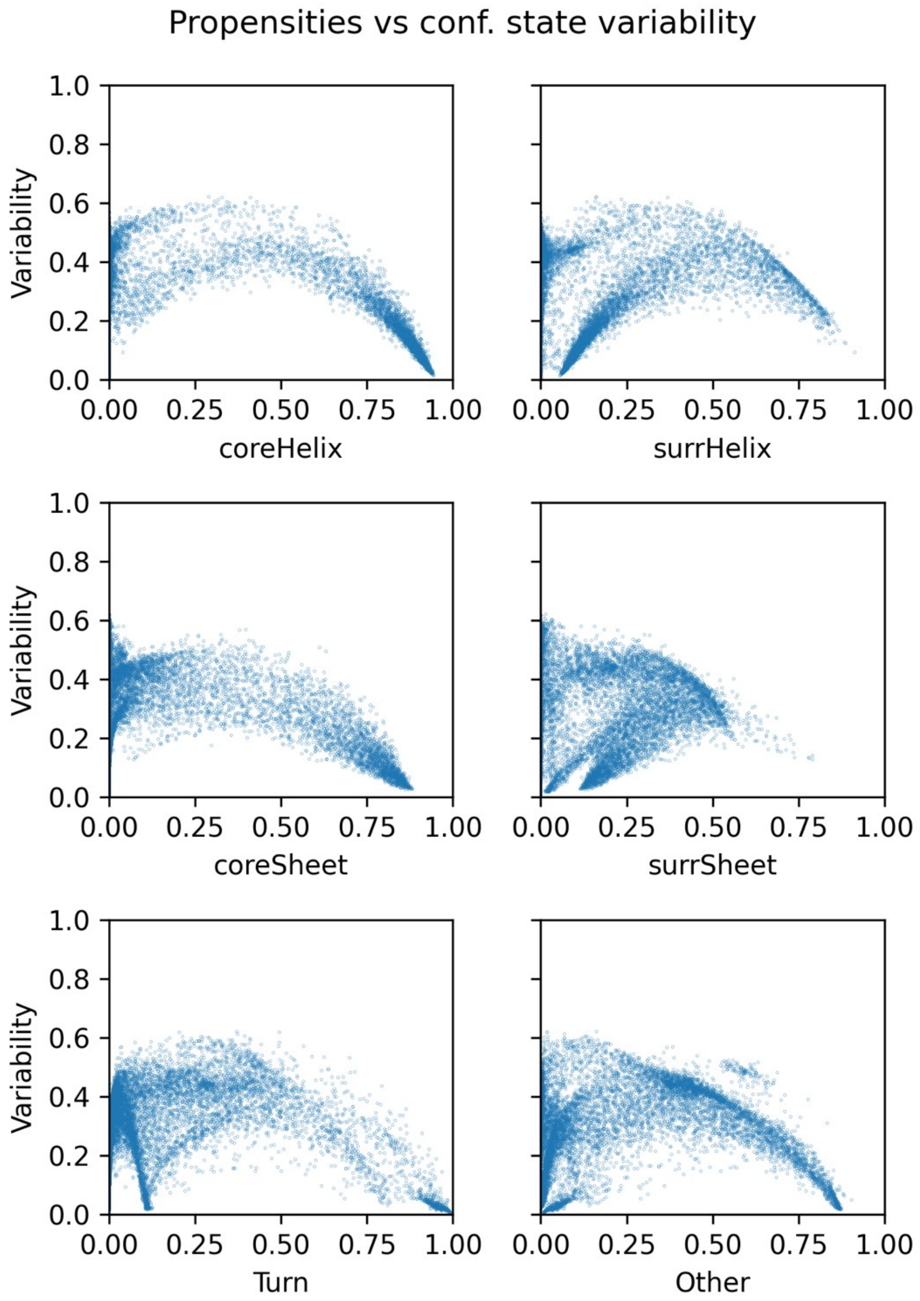

***Supplementary Fig. 13. Conformational states propensities vs conformational state variability.*** *The metrics were calculated using bootstrapping with a sample size of 3 (10,000 samples). Conformational state propensities for each conformational state are plotted on the x axes, while the y axes show the conformational state variability. As the conformational state propensity for a single conformational state approaches 1, the conformational state variability drops to 0. In contrast, no such correlation can be observed in the left half of the plots. As the probability for a given conformational state approaches 0, the residue may still assume any of the remaining five states, and transition between them, thus, allowing for no conclusions on the conformational state variability.*

This work is shared under a CC BY 4.0 license.

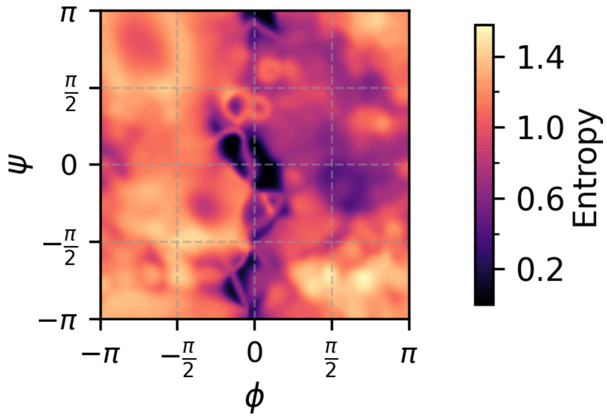

***Supplementary Fig. 14. Entropy of the conformational states in the (ϕ, ψ)-space.*** *The entropy is a measure of how sensitive the conformational states are to minor changes in the backbone dihedrals in the corresponding areas. The area around (0, 0) is not populated in any of the conformational state models. Thus, the entropy in this region approaches 0.*

This work is shared under a CC BY 4.0 license.

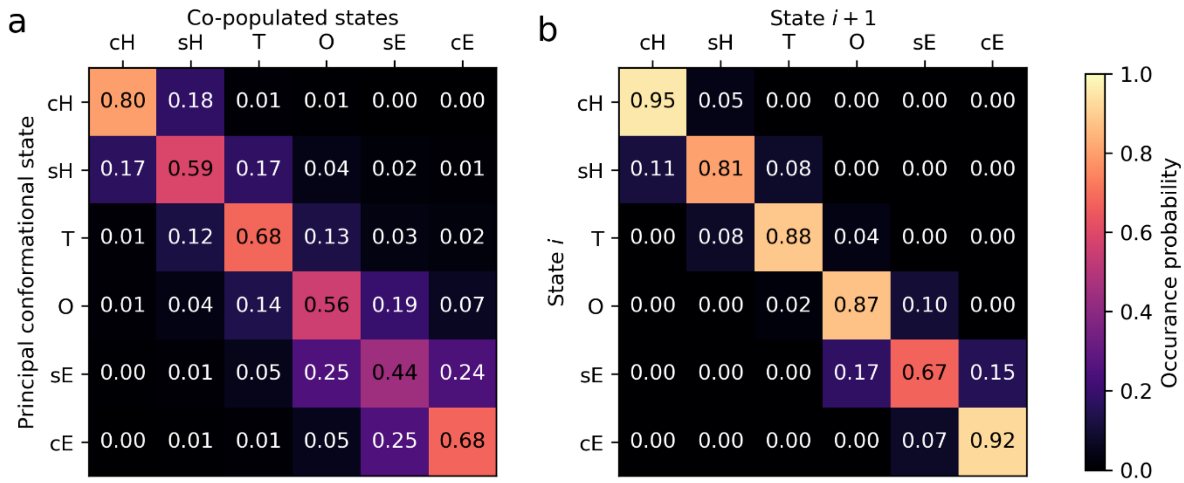

***Supplementary Fig. 15. Co-occurrence of conformational states.*** *a) This plot shows the overall co-occurrence of conformational states in individual residues. For every residue with a given most likely conformational state (vertical axis), the average remaining propensities for all other states are shown (horizontal axis). b) This plot shows the probabilities of residues transitioning between conformational states between two subsequent time points. Both plots were obtained with window size 3. The labels correspond to the conformational states as follows: cH* (Core helix)*, sH (*Surrounding helix*), T (*Turn*), O* (Other)*, sE (*Surrounding sheet*), cE (*Core sheet*).*

This work is shared under a CC BY 4.0 license.

### Supplementary Data

#### Supplementary Data 1: **List of proteins according to order class**

**Disordered:** 2aqc, 1xq8, 1mjc, 1u96, 1ss3, 2kbz, 2l95, 1nq4, 1vd0, 1y00, 1rk9, 2l50, 1yob, 1rqs, 1czh

**Half-disordered:** 2jo6, 2rvh, 2khz, 1akp, 2m2u, 2ho9, 2m9v, 2joy, 2mlw, 2lnu, 1b75, 2ot2, 1c9h, 2l7w, 1r1t, 2l49, 2l9n, 2myi, 2ls7, 1o13, 2lvh, 2l6o, 5vo7, 2kly, 2l25, 1wm2, 2kpq, 2kg7, 1l7y, 2gqb, 1eni, 1z2m, 2den, 2jra, 2jyz, 2jz4, 2m0y, 2m2a, 2mc9, 2mhe, 2mvz, 2x8n, 4beh, 4eyv, 5jtk

**Structured:** 2mkx, 1jqr, 2fm4, 1jw3, 2rr6, 1hg7, 1sp0, 2kva, 3hnx, 1tm9, 1om2, 2d49, 2hfi, 1ufn, 1qkh, 2fvt, 1rzs, 1x3a, 1ey1, 1wjt, 2hgk, 2k2t, 2jny, 2jso, 2rn4, 1bq0, 1g9l, 1imq, 1iyr, 1jh3, 1jw2, 1mp1, 1nnv, 1nzp, 1rq6, 1rrz, 1ryk, 1t0g, 1tl6, 1v66, 1v9v, 1xhs, 2ebm, 2end, 2gbs, 2hfq, 2jgx, 2jov, 2jrr, 2jv2, 2jzv, 2ken, 2lfj

#### Supplementary Data 2: **List of proteins mapped to NMR metrics**

Only the following subset of MD proteins had their corresponding Random Coil Index information available:

2m2a, 1jw2, 2ls7, 1t0g, 2jny, 1tm9, 1v66, 2gqb, 2ebm, 2kpq, 1ryk, 2gbs, 2fvt, 2ken, 1b75, 2l25, 2rr6, 1iyr, 2hfi, 2ho9, 1akp, 2fm4, 1xhs, 1l7y, 1ufn, 2mlw, 1ey1, 2kva, 2mvz, 2lnu, 2lfj, 1om2, 2jyz, 2mhe, 1v9v, 2l49, 1jqr, 1x3a, 1g9l, 2rvh, 5vo7, 1bq0, 2l6o, 1c9h, 2jov, 2ot2, 2joy, 2rn4, 2hgk, 1jh3, 2jz4, 5jtk, 1rzs, 2jso, 2kbz, 1rq6, 2jzv, 1wjt, 1jw3, 2jrr, 1hg7, 1imq

This work is shared under a CC BY 4.0 license.
